# Dynamics of gene expression in single root cells of *A. thaliana*

**DOI:** 10.1101/448514

**Authors:** Ken Jean-Baptiste, José L. McFaline-Figueroa, Cristina M. Alexandre, Michael W. Dorrity, Lauren Saunders, Kerry L. Bubb, Cole Trapnell, Stanley Fields, Christine Queitsch, Josh T. Cuperus

**Affiliations:** Department of Genome Sciences, University of Washington, Seattle, WA 98195.; Department of Biology, University of Washington, Seattle, WA 98195.; Department of Medicine, University of Washington, Seattle, WA 98195.

**Author notes:** The author responsible for distribution of materials integral to the findings presented in this article in accordance with the policy described in the Instructions for Authors (www.plantcell.org) is: Josh Cuperus.

## Abstract

Single-cell RNA-seq can yield high-resolution cell-type-specific expression signatures that reveal new cell types and the developmental trajectories of cell lineages. Here, we apply this approach to *A. thaliana* root cells to capture gene expression in 3,121 root cells. We analyze these data with Monocle 3, which orders single cell transcriptomes in an unsupervised manner and uses machine learning to reconstruct single-cell developmental trajectories along pseudotime. We identify hundreds of genes with cell-type-specific expression, with pseudotime analysis of several cell lineages revealing both known and novel genes that are expressed along a developmental trajectory. We identify transcription factor motifs that are enriched in early and late cells, together with the corresponding candidate transcription factors that likely drive the observed expression patterns. We assess and interpret changes in total RNA expression along developmental trajectories and show that trajectory branch points mark developmental decisions. Finally, by applying heat stress to whole seedlings, we address the longstanding question of possible heterogeneity among cell types in the response to an abiotic stress. Although the response of canonical heat shock genes dominates expression across cell types, subtle but significant differences in other genes can be detected among cell types. Taken together, our results demonstrate that single-cell transcriptomics holds promise for studying plant development and plant physiology with unprecedented resolution.

## INTRODUCTION

Many features of plant organs such as roots are traceable to specialized cell lineages and their progenitors (Irish, 1991; Petricka et al., 2012). The developmental trajectories of these lineages have been based on tissue-specific and cell-type-specific expression data derived from tissue dissection and reporter gene-enabled cell sorting (Birnbaum et al., 2003; Brady et al., 2007; Li et al., 2016). However, tissue dissection is labor-intensive and imprecise, and cell sorting requires prior knowledge of cell-type-specific promoters and genetic manipulation to generate reporter lines. Few such lines are available for plants other than the reference plant *Arabidopsis thaliana* (Rogers and Benfey, 2015). Advances in single-cell transcriptomics can replace these labor-intensive approaches. Single-cell RNA-seq has been applied to heterogeneous samples of human, worm, and virus origin, among others, yielding an unprecedented depth of cell-type-specific information (Cao et al., 2017; Irish, 1991; Packer and Trapnell, 2018; Russell et al., 2018; Trapnell, 2015; Trapnell et al., 2014).

While several examples of single cell RNA-seq have been carried out in *Arabidopsis* (Efroni et al., 2016, 2015; Brennecke et al., 2013), they were restricted to only a few cells or cell types. No whole organ single-cell RNA-seq has been attempted in any plant species. The *Arabidopsis* examples focused on root tips, finely dissecting the dynamics of regeneration or assaying technical noise across single cells in a single cell type. Thus, a need exists for larger scale technology that allows a more complete characterization of the dynamics of development across many cell types in an unbiased way. Such technology would increase our ability to assay cell types without reporter gene-enabled cell sorting, identify developmental trajectories, and provide a comparison of how different cell types respond to stresses or drugs. Several high-throughput methods have been described for sequencing of RNA at a high throughput of single cells. Most of these, including most droplet-based methods, rely on the 3’ end capture of RNAs. However, unlike with bulk RNA-seq, the data from single cell methods can be sparse, such that genes with low expression can be more difficult to study. Here, we take advantage of expression data from root-specific reporter lines in *A. thaliana* (Birnbaum et al., 2003; Brady et al., 2007; Cartwright et al., 2009; Li et al., 2016) to explore the potential of single-cell RNA-seq to capture the expression of known cell-type-specific genes and to identify new ones. We focus on roots of mature seedlings and probe the developmental trajectories of several cell lineages.

## RESULTS

### Single cell RNA-seq of whole A. thaliana roots reveals distinct populations of cortex, endodermis, hair, non-hair, and stele cells

We used whole *A. thaliana* roots from seven-day-old seedlings to generate protoplasts for transcriptome analysis using the 10x Genomics platform (**Supplemental Figure 1A**). We captured 3,121 root cells to obtain a median of 6,152 unique molecular identifiers (UMIs) per cell. UMIs here are 10 base random tags added to the cDNA molecules that allow us to differentiate unique cDNAs from PCR duplicates. These UMIs corresponded to the expression of a median of 2,445 genes per cell and a total of 22,419 genes, close to the gene content of *A. thaliana*. Quality measures for sequencing and read mapping were high. Of the approximately 79,483,000 reads, 73.5% mapped to the TAIR10 *A. thaliana* genome assembly, with 67% of these annotated transcripts. These values are well within the range reported for droplet-based single-cell RNA-seq in animals and humans.

For data analysis, we applied Monocle 3, which orders transcriptome profiles of single cells in an unsupervised manner without *a priori* knowledge of marker genes (Qiu et al., 2017a; Qiu et al., 2017b; Trapnell et al., 2014). We used the 1500 genes in the data set (**Supplemental Data Set 1**) that showed the highest variation in expression (**Supplemental Figure 1B**). For unsupervised clustering, we used 25 principal components (PC). These 25 PCs accounted for 72.5% of the variance explained by the first 100 PCs, with the first PC explaining 11% and the 25th PC explaining 0.9% (**Supplemental Figure 1C**). Cells were projected onto two dimensions using the uniform manifold approximation and projection (UMAP) method (McInnes and Healy, 2018) and clustered, resulting in 11 clusters (**Figure 1A**) (Blondel et al., 2008). Most clusters showed similar levels of total nuclear mRNA, although clusters 9 and 11 were exceptions with higher levels (**Supplemental Figure 1D**). Because some of the UMAP clusters, specifically clusters 9 and 11, consisted of cells that had higher than average amounts of nuclear mRNA, we were concerned that these clusters consisted merely of cells that were doublets, *i.e.* two (or more) cells that received the same barcode and that resulted in a hybrid transcriptome. As cells were physically separated by digestion, it was possible that two cells remained partially attached. In order to identify potential doublets in our data, we performed a doublet analysis using Scrublet (Wolock et al., 2018), which uses barcode and UMI information to calculate the probability that a cell is a doublet. This analysis identified only 6 cells, of 3,021 cells analyzed, as doublets, spread across multiple UMAP clusters and multiple cell types (**Supplemental Figure 1E**). Overall, given the low number of doublets, we did not attempt to remove these cells.

**Figure 1.**
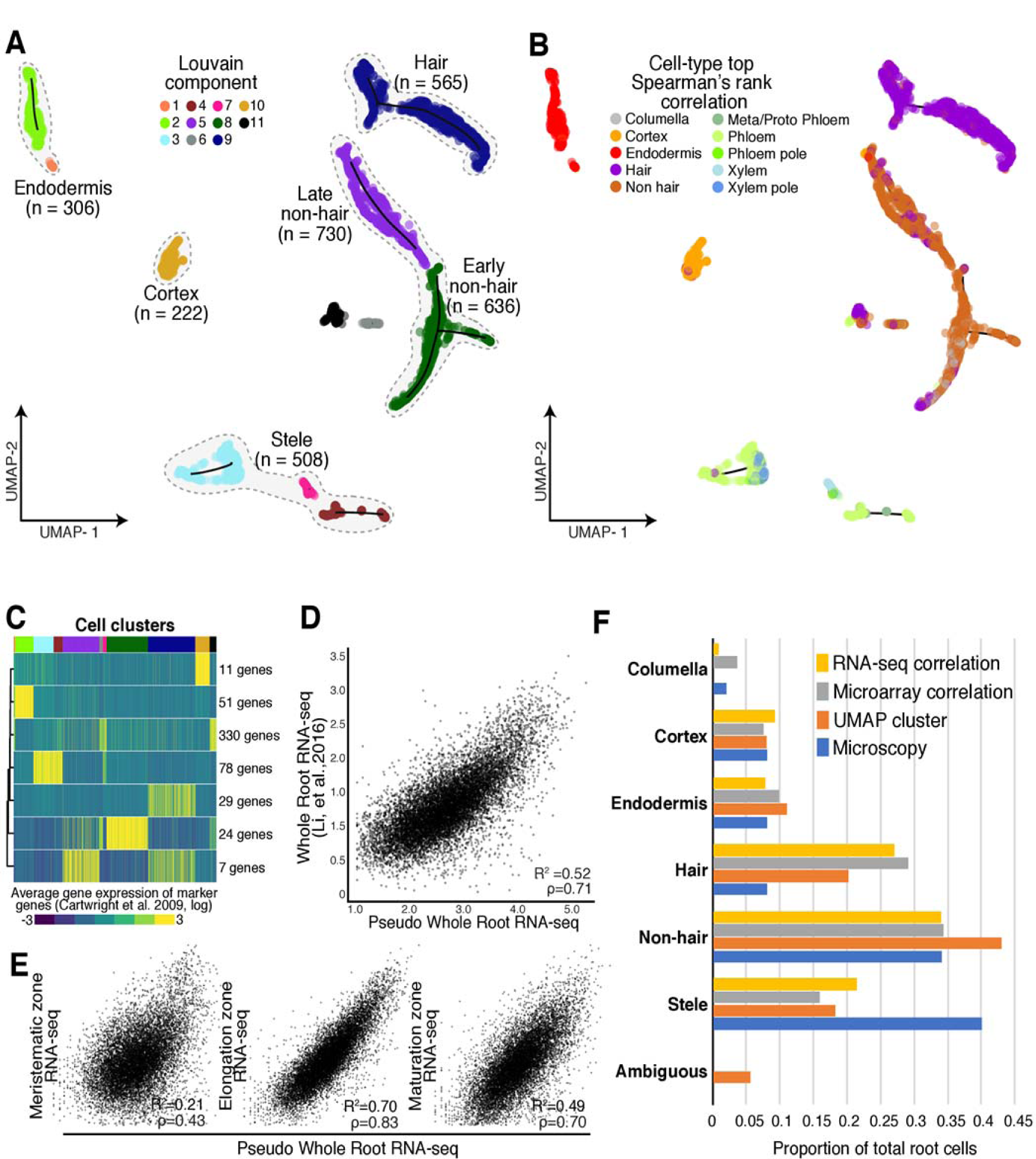
Annotation of cell and tissue types for single-cell RNA-seq of whole A. thaliana roots. **(A)** Root cells were clustered and projected onto two-dimensional space with UMAP (McInnes and Healy, 2018) Filled circles represent individual cells; colors represent their respective Louvain component. Monocle 3 trajectories (black lines) are shown for clusters in which a trajectory could be identified. **(B)** Riled circles represent individual cells; colors indicate cell and tissue type based on highest Spearman’s rank correlation with sorted tissue-specific bulk expression data (Brady et al., 2007; Cart-wright et al. 2009)‘ **(C)** Known marker genes (Brady et aL, 2007; Cartwright et al., 2009) were used to ciuster single-cell gene expression profiles based on similarity. The expression of 530 known marker genes was grouped into 7 clusters, using k-means clustering. Mean expression for each cluster (rows) is presented for each cell (columns). Cells were ordered by their respective Louvain component indicated above by color (see A, starting at component 1 at left). Number of genes in each cluster is denoted at right. **(D)** Single-cell RNA-seq pseudo-bulked expression data are compared to bulk expression data 01 whole roots (Li et 31., 2016). **(E)** Single-cell pseudo-bulk expression data are compared to bulk expression data of the three developmental regions of the A. thaliana root (Li et al., 2016). **(F)** Proportions oi cells as annotated by either UMAP (in A), Spearman’s rank correlation (in B), or Pearson’s rank in Supplemental Figure 2, are compared to proportions determined by microscopy (Brady et al., 2007; Cartwright et al., 2009).

To assign these clusters to cell types, we performed three complementary analyses relying on two expression data sets from tissue-specific and cell-type-specific reporter lines: an earlier one generated with microarrays (Brady et al., 2007; Cartwright et al., 2009) and a more recent one generated with RNA-seq and a greater number of lines (Li et al., 2016). We first compared the microarray expression data for each reporter line to the gene expression values in each single cell, using Spearman’s rank correlations to assign each cell a cell type identity based on highest correlation of gene expression (**Figure 1B, Supplemental Data Set 2**) (Brady et al., 2007; Cartwright et al., 2009). Second, we compared the RNA-seq expression data to the gene expression values in each single cell by Pearson’s correlation (Li et al., 2016, **Supplemental Figure 2A**). Third, we examined the expression of 530 cell-type-specific marker genes (Brady et al., 2007) by defining seven marker gene clusters with k-means clustering and calculating their average expression for each cell. We then compared each cell’s UMAP Louvain component cluster assignment (**Figure 1A**) with its marker-gene-based assignment. Louvain components were derived using the Louvain method for community detection (Blondel et al., 2008) which is implemented in Monocle 3. Unlike k-means clustering for which the user provides the desired number of clusters to partition a dataset, Louvain clustering optimizes modularity (*i.e.* the separation of clusters based on similarity within a cluster and among clusters), aiming for high density of cells within a cluster compared to sparse density for cells belonging to different clusters. The 11 clusters presented in **Figure 1A** optimized the modularity of the generated expression data and were not defined by us.

In general, the UMAP clusters showed high and cluster-specific expression of marker genes. For example, cells in cluster 10 showed high and specific mean expression of cortex marker genes (**Figure 1C, Supplemental Figure 3, Supplemental Data Set 3**). Both expression correlations and marker gene expression allowed us to assign the Louvain components to five major groups: root hair cells, non-hair cells (containing both an early and late cluster), cortex cells, endodermis cells and stele cells (containing both xylem and phloem cells) (**Figure 1A**). Although some cells were most highly correlated in expression with the cell type columella in Spearman’s rank tests and RNA-seq Pearson’s correlation, these cells co-clustered with non-hair cells (**Figure 1B, Supplemental Figure 2**). This finding is consistent with bulk RNA-seq data of sorted cells (Li et al., 2016). Specifically, the *PET11* (columella)-sorted bulk RNA-seq data are most similar to bulk RNA-seq data sorted for *GL2* and *WER* (Li et al., 2016), both of which mark non-hair cells (Petricka et al., 2012). Therefore, these cells were grouped as early non-hair cells with other non-hair cells in Louvain component 8. As their expression values were best correlated with RNA-seq data for *WER*-sorted cells, they likely represent a mix of early non-hair and lateral root cap cells, which have very similar expression profiles (**Supplemental Figure 2**).

We assessed the extent to which combined single-cell root expression data resembled bulk whole root expression data (Li et al., 2016) (**Figure 1D, E**). We observed strong correlations between these two data sets (Pearson’s correlation coefficient [R^2^] = 0.52, Spearman’s ρ=0.71). We also compared the combined single-cell expression data to three bulk expression data sets representing the major developmental zones in the *A. thaliana* root: the meristematic zone, the elongation zone, and the maturation zone (**Figure 1E**). We observed the highest correlation of single-cell and bulk expression in the elongation zone (R^2^=0.70, ρ=0.83) and a lower correlation in the maturation zone (R^2^=0.58, ρ=0.70). This observation is surprising given the more mature developmental stage of the harvested roots (**Supplemental Figure 1A**), and likely reflects that younger cells are more easily digested during protoplasting and contribute in greater numbers to the gene expression data. As expected, single-cell and bulk expression were poorly correlated in the meristematic zone (R^2^=0.11, ρ=0.43), as meristematic tissue accounts for only a small proportion of mature roots. Furthermore, we compared tissue-specific expression (Li et al., 2016) to expression both in the annotated cell clusters and in cells expressing appropriate marker genes. In general, we found strong correlations among these data sets, suggesting that the clusters are annotated correctly (**Supplemental Table 1**).

We also compared the relative representation of root cell types between our data set and estimates based on microscopy studies (**Figure 1F**) (Brady et al., 2007; Cartwright et al., 2009). Independent of annotation method, we observed the expected numbers of cortex (222 Spearman’s/ 233 UMAP), endodermis (306/ 304), non-hair cells (1201/ 1061) and columella cells (111/ no UMAP cluster). Hair cells (565/ 898) were overrepresented whereas stele cells (508/ 490) were underrepresented, possibly reflecting a bias in the protoplast preparation of whole root tissue.

Protoplasting, the removal of the plant cell wall, alters the expression of 346 genes (Birnbaum et al., 2003); 76 of these genes were included in the 1500 genes with the highest variation in expression (**Supplemental Data Set 1, Supplemental Figure 1B**) that we used for clustering. Some of the 76 genes showed cell-type-specific expression. To exclude the possibility that the expression pattern of these genes produced artefactual clusters and cell-type annotations, we removed them from the analysis and re-clustered, which resulted in a similar UMAP visualization, with similar numbers of Louvain components and cell types.

### Single-cell RNA-seq of identifies novel genes with cell-type and tissue-type-specific expression

Some marker genes are not expressed exclusively in a single cell type, making it desirable to identify additional genes with cell-type-specific expression. We first confirmed the high and cluster-specific expression of well-known marker genes (**Figure 2A, Supplemental Figure 4**) (Li et al., 2016) such as the root-hair-specific *COBL9*, the endodermis-specific *SCR* and the three stele-specific genes *MYB46* (xylem-specific), *APL* (phloem-specific), and *SUC2* (phloem-specific). The nonspecific expression of the quiescent center cell marker genes *WOX5* and *AGL42* is likely due to the failure to capture sufficient numbers of these rare cells. The nonspecific expression of *WOL* and the more heterogeneous pattern of both *WER* and *GL2* expression have been previously observed (Brady et al., 2007; Winter et al., 2007). Second, to find novel marker genes, we identified genes with significantly different expression within and among Louvain component clusters by applying the Moran’s I test implemented in Monocle 3. We found 317 genes with cluster-specific expression, 164 of which were novel, including at least one in each cluster (**Figure 2A, Supplemental Data Set 4)**. Using cell-type annotations rather than Louvain clusters, we identified 510 genes with cell-type-specific expression, of which 317 overlapped with the Louvain component cluster-specific expression genes, as well as an additional 125 novel genes, some of which have been implicated in the development of a cell lineage in targeted molecular genetics studies.

**Figure 2.**
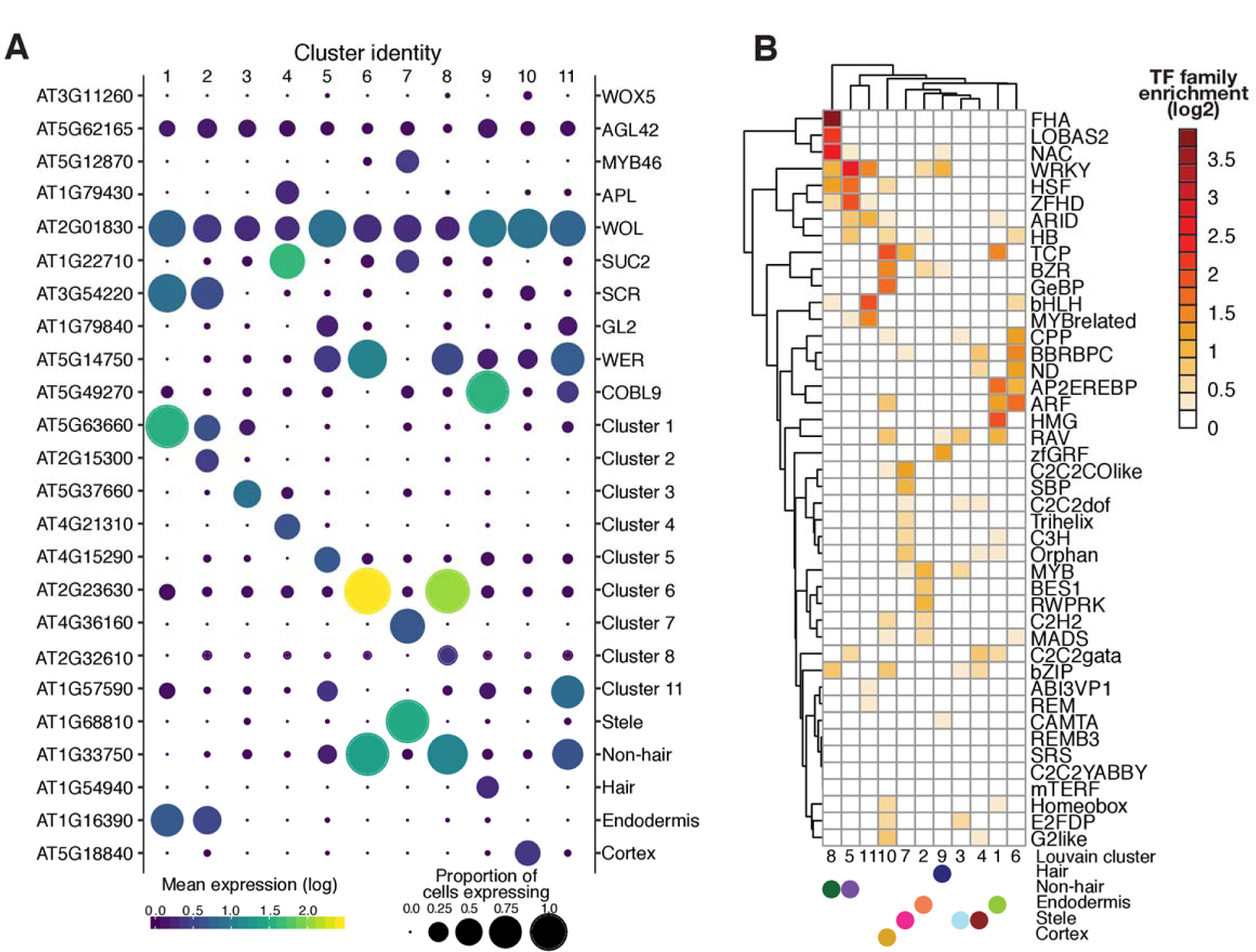
Novel cluster-specific and tissue-specific genes and enriched transcription factor motifs. **(A)** Proportion of cells (circle size) and mean expression (circle color) of genes with cluster-specific and tissue-specific expression are shown, beginning with known marker genes labeled with their common name (right) and their systematic name (left). For novel genes, the top significant cluster-specific genes are shown, followed by the top significant tissue-specific genes; both were identified by principal graph tests (Moran’s I) as implemented in Monocle 3. Note the correspondence between Louvain components and cell and tissue types. For all novel cluster-specific and tissue-specific genes, see Supplemental Table 3. **(B)** Enrichments of known transcription factor motifs (O’Malley et al., 2016) 500 bp upstream of genes with ckjster-spedfic expression compared to genome background. Motifs are specific to transcription factor gene families rather than individual genes. The plot is clustered based an similarity in enrichments with Louvain components and cell and tissue types (filled circles) indicated.

For example, the stele-specific AT1G8810 (*ABS5, T5L1*) gene (cluster 7, **Figure 2A**) encodes a bHLH protein that promotes vascular cell division and differentiation as part of a heterodimer with second bHLH protein, LHW (Katayama et al., 2015; Ohashi-Ito et al., 2014). Another stele-specific gene, AT4G36160 (*ANAC076*, *VND2*) (cluster 7), encodes a ClassIIB NAC-domain transcription factor that contributes to xylem vessel element differentiation by promoting secondary cell wall formation and programmed cell death (Tan et al., 2018). In tissue-specific bulk data (Brady et al., 2007; Winter et al., 2007), both genes show xylem-specific expression consistent with their biological functions; *T5L1* expression is high only in the meristematic and elongation zones, while *VND2* expression starts in the elongation zone and persists throughout the maturation zone. Other genes, not previously implicated in root development, show tissue-specific bulk expression patterns consistent with the single-cell data. For example, AT1G54940 (*GUX4*, *PGSIP4*), which encodes a xylan glucuronosyltransferase (Lee et al., 2012; Mortimer et al., 2010), was specifically expressed in hair cells (cluster 9) and is most highly expressed in cells destined to become hair cells in the elongation zone and in differentiated hair cells in the maturation zone (Brady et al., 2007; Cartwright et al., 2009).

### Expression of some transcription factors shows high correlation with specific cell types

We asked whether we could identify transcription factors that may contribute to the cluster-specific expression patterns. To do so, we tested for transcription factor motif enrichments in the proximal regulatory regions of genes with cluster-specific expression, examining 500 bp upstream of the transcription start site (Alexandre et al., 2018; Sullivan et al., 2014) and a comprehensive collection of *A. thaliana* transcription factor motifs (O’Malley et al., 2016). This analysis revealed significant transcription factor motif enrichments among clusters and annotated major tissues and cell types (**Figure 2B**).

As transcription factors in *A. thaliana* often belong to large gene families without factor-specific motif information (Riechmann et al., 2000), it is challenging to deduce the identity of the specific transcription factor that drives cluster-specific transcription factor motif enrichment and expression. As an approximation, we examined transcription factors that were expressed in the cluster or tissue in which a significant enrichment of their motif was found, or in neighboring cell layers (some factors move between cells (Petricka et al., 2012)) (**Supplemental Data Set 4**). We focused first on the small *BZR/BEH* gene family whose motif was specifically enriched in cortex cells (cluster 10). Of the six genes (BEH1/AT3G50750, BEH2/AT4G36780, BEH3/AT4G18890, BEH4/AT1G78700, BES1/AT1G19350, and BZR1/AT1G75080) the single recessive *beh4*, *bes1*, and *bzr1* mutants exhibit altered hypocotyl length (Lachowiec et al., 2018). Double mutant analysis suggests partial functional redundancy, which agrees with our observation of overlapping expression patterns for these genes across cell types (**Supplemental Figure 5A, B**). In contrast, neither *beh1* and *beh2* single mutants nor the respective double mutant show phenotypic defects (Lachowiec et al., 2018). However, *BEH2* was the most highly expressed *BZR/BEH* family member across clusters and annotated root tissue and cell types (**Supplemental Figure 5A, B**). Although *BEH4*, the most ancient family member with the strongest phenotypic impact, showed cortex-specific expression, none of the *BZR/BEH* genes showed significance for cluster-specific expression, suggesting that combinations of family members, possibly as heterodimers, may result in the corresponding motif enrichment in cortex cells (**Supplemental Figure 5A, B**). In particular, *BES1* and *BZR* expression was highly correlated, consistent with these genes being the most recent duplicates in the family (**Supplemental Figure 5C**) (Lachowiec et al., 2013; Lan and Pritchard, 2016).

In contrast to the *BEH/BZR* gene family, we found stronger cluster specificity for some TCP transcription factors. The TCP motif was strongly enriched in cortex (cluster 10), endodermis (cluster 1) and stele (cluster 7). Of the 24 TCP transcription factors, we detected expression for eight. Of these, *TCP14* (AT3G47620) and *TCP15* (AT1G69690) were expressed primarily in stele (clusters 7 and 4) although this cluster-specific expression was not statistically significant (**Figure 2B, Supplemental Figure 5D, E, Supplemental Data Set 4**). *TCP14* and *TCP15* are class I TCP factors thought to promote development. Acting together, *TCP14* and *TCP15* promote cell division in young internodes (Kieffer et al., 2011), seed germination (Resentini et al., 2015), cytokinin and auxin responses during gynoecium development (Lucero et al., 2015), and repression of endoreduplication (Peng et al., 2015). Both genes are expressed in stele in bulk tissue data (Brady et al., 2007; Winter et al., 2007), with *TCP14* expression also observed in the vasculature by *in situ* hybridization (Tatematsu et al., 2008). *TCP14* can affect gene expression in a non-cell-autonomous manner.

To further investigate the co-occurrence of cluster-specific transcription factor motif enrichments with transcription factor expression, we next examined the novel genes with significant cluster-specific expression. Eight of these encode transcription factors with corresponding highly enriched cluster-specific binding motifs. For one of these, *BRN2* (AT4G10350), cluster-specific expression coincided with enrichment of the NAC transcription factor family motif(cluster 8, non-hair and lateral root cap cells, **Figure 2B**). *BRN2* encodes a ClassIIB NAC transcription factor implicated in root cap maturation together with *BRN1* and *SMB*. Class IIB NAC transcription factors are thought to contribute to terminal cell differentiation accompanied by strong cell wall modifications (Bennett et al., 2010). In our data, *BRN2* was most highly expressed in cluster 8 (non-hair and lateral root cap cells) and less so in cluster 6 (**Supplemental Data Set 4**).

### Clustering stele cells identifies novel genes with cell-type specific expression in the vasculature

Our initial attempts to annotate and separate cell types within stele tissue with marker gene expression or Spearman’s rank correlations failed. Instead, we separately clustered stele cells to reveal 6 sub-clusters upon UMAP visualization, with 5 sub-clusters containing more than 40 cells. Their annotation via Spearman’s rank correlation with sorted bulk data was not successful; however, using well-established marker genes expression, we detected cluster-specific expression patterns (**Figure 3A and B**).

**Figure 3.**
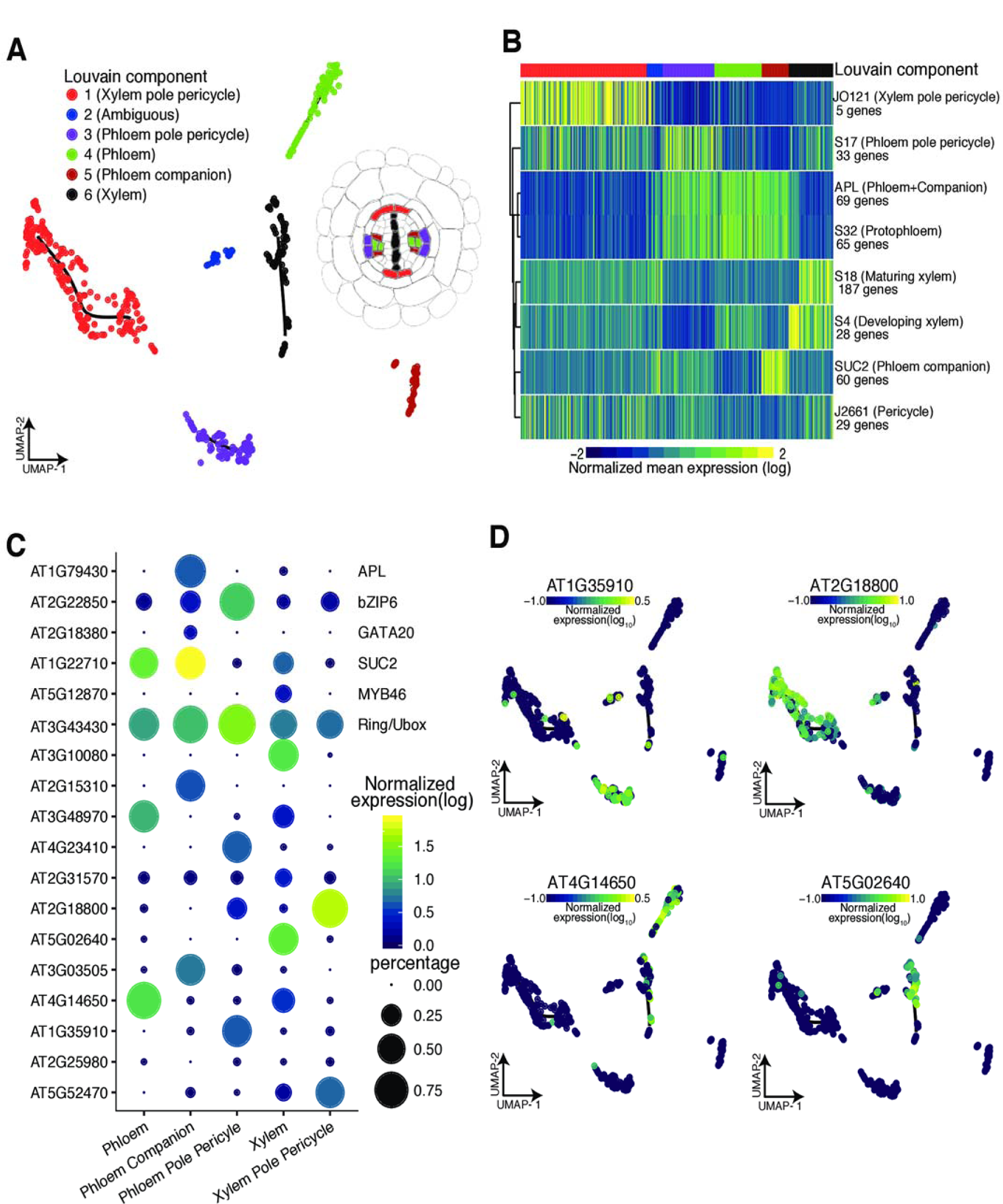
Re-clustering of stele cells yields distinct sub-clusters of vasculature cell types. **(A)** Cells initially annotated as stele tissue were re-clust8red, resulting in six distinct sub-clusters cells, five of which contained more than 40 cells, **(B)** Mean expression for previously identified cell-type specific genes (Cartwright et al., 2009) in each cell is shown, allowing annotation of stele sub-cluster identities as shown in (A). **(C)** Proportion of cells (circle size) and mean expression (circle color) 01 genes with cluster-specific and tissue-specific expression are shown, starting with known marker genes at the top, labeled with their common name (right) and their systematic name (left). Below, novel significant tissue-specific genes are shown with their systematic names, identified by principal graph tests (Moran’s I) as implemented in Monocle 3. **(D)** Example expression overlays for cluster-specific genes identified by the principal graph test in (C).

Cells closely related to xylem pole pericycle constituted the largest group of cells (205 cells); phloem pole pericycle cells were the second largest (84 cells). The high number of pericycle cells likely reflects our experimental procedure, as these cells reside on the exterior of the vascular bundle. Both phloem and xylem clusters showed similar numbers of cells (77 cells and 72 cells respectively); the phloem companion cells formed a distinct cluster. We observed the expected sub-cluster expression for several known genes and marker genes and identified novel genes with sub-cluster-specific expression (**Figure 3C, D, Supplemental Data Set 1**). Although there was some discrepancy, especially for the *APL* gene, which is expressed in both companion and phloem cells (**Figure 3C)**, this is largely due to missing data.

### Pseudotime trajectories coincide with the development stages of cortex, endodermis, and hair cells

We next sought to visualize the continuous program of gene expression changes that occurs as each cell type in root differentiates. Because whole roots contain a mix of cells at varying developmental stages, we reasoned that our experiment should have captured a representative snapshot of their differentiation. Monocle not only clusters cells by type but also places them in “pseudotime” order along a trajectory that describes their maturity. To make these trajectories, Monocle 3 learns an explicit principal graph from the single-cell-expression data through reversed graph embedding, an advanced machine learning method (Qiu et al., 2017a; Qiu et al., 2017b; Trapnell et al., 2014). To dissect the developmental dynamics of individual clusters, we first focused on the well-defined root-hair cells, in which combined single-cell expression values highly correlated with those from bulk protoplasts sorted for expression of the *COBL9* root-hair marker gene (**Supplemental Table 1**). To annotate the unsupervised trajectory that Monocle 3 created for hair cells, we used the Spearman’s rank test to compare expression in all cells to bulk expression data representing 13 different developmental stages in root tissues from all the available sorted cell types (**Supplemental Figure 6**) (Brady et al., 2007; Cartwright et al., 2009). Each cell was assigned the developmental stage and cell type most correlated with its expression values (**Figure 4A)**. The hair cells with the earliest developmental stage assignment were designated as the root of the trajectory. Next, pseudotime was calculated for all other hair cells based on their distance from the root of the trajectory (**Figure 4B**). We compared this calculated pseudotime with the most highly correlated developmental assignment from bulk data, finding close agreement (**Figure 4B**). Examples of genes that are expressed early and late in pseudotime in the UMAP hair cluster are shown in **Figure 4C**.

**Figure 4.**
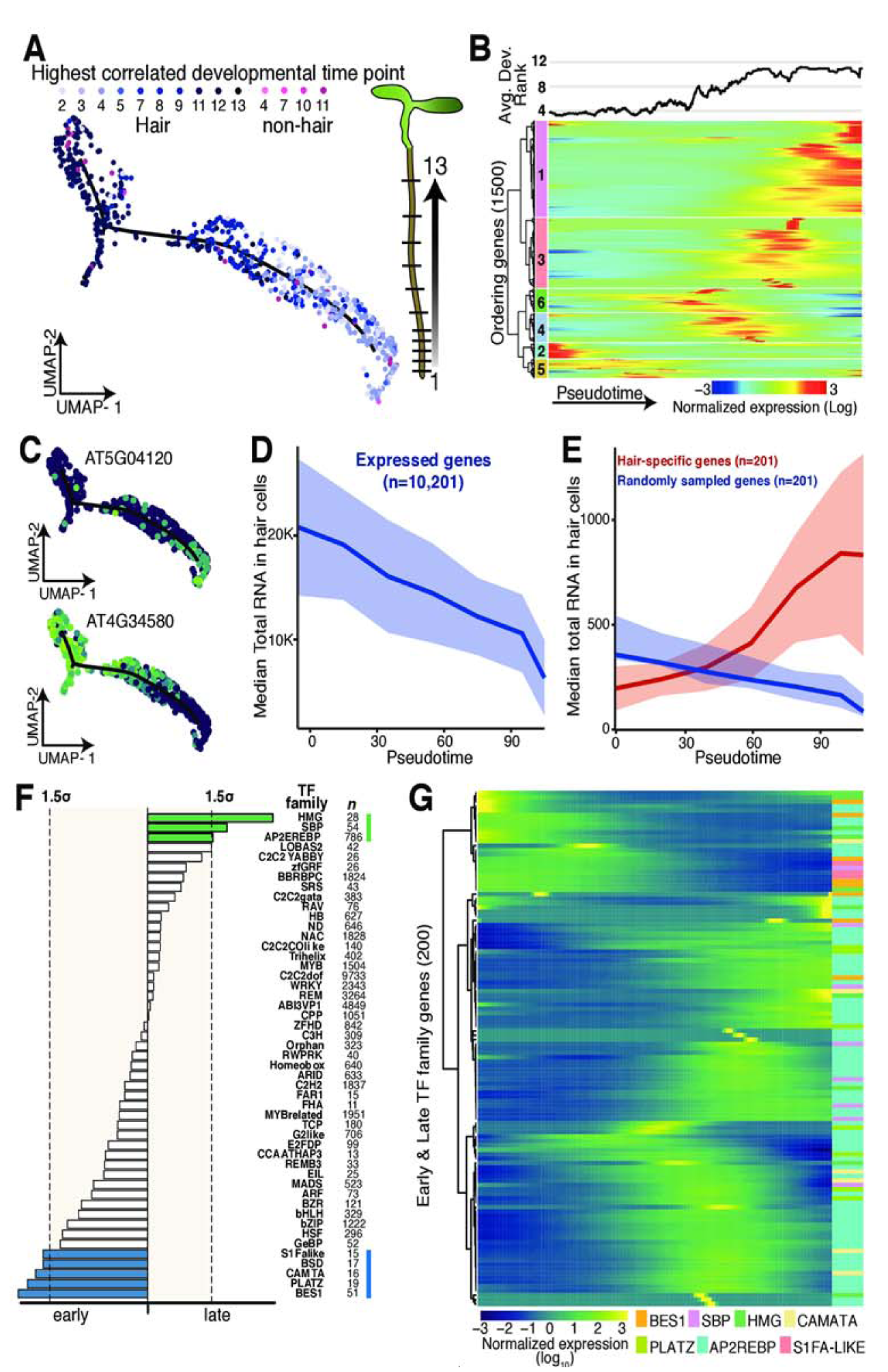
Developmental trajectory of hair cells. **(A)** UMAP-clustered hair cells were assigned a developmental time point based on highest Spearman’s rank correlation with bulk expression data of staged tissue (13 developmental stages) (Brady et al., 2007; Cartwright et al., 2009). Cell type and developmental time points are indicated in shades of blue (and pink). Graphic illustrates developmental stages in A, thaliana root (Plant lllusirations). **(B)** Cells were ordered in pseudotime; columns represent cells, rows represent expression of the 1500 ordering genes. Rows were grouped based on similarity in gene expression, resulting iri 6 clusters (indicated left), with genes in clusters 2 and 5 expressed early in pseudotime and genes in cluster 1 expressed late. Hair cells with the earliest developmental signal (Brady et al., 2007; Cartwright et al, 2009) were designated as the root oí the trajectory. The graph above represents the aver-age best-correiation of developmental stage (Brady et al., 2007; Cartwright et al., 2009) in a scrolling window of 20 cells with pseudotime, showing the expected increase in developmental age with increasing pseudotime, **(C)** Examples of an early and a late expressed hair-cell-specific qene. Gene expression in each cell is superimposed onto the UMAP cluster and trajectorv, with lighter colors indicating higher gene expression **(D)** Median total RNA captured in cells decreases across pseudotime. Number 0! genes included is indicated. **(E)** Comparison of median total RNA for hair-cell-specific genes (in red) to a comparable random set of genes (in blue). Number of genes is indicated (Permutation test p-value ~ 1 × 10-4). **(F)** Different transcription factor motifs reside in the 500 bp upstream regions of genes expressed early (clusters 2,5) compared to genes expressed late (cluster 1). Transcription factor motifs specific to early hair cells are denoted with blue bars, those for late hair cells with green bars; bar length indicates motif frequency. Thresholds on either side (grey box, dotted lines) refer ίο 1.5 standard deviation above mean motif frequency. **(G)** Expression of individual members of transcription factors families highlighted in D across pseudotime identifies candidate factors driving early or late gene expression.

Hair cells undergo endoreduplication as they mature, resulting in up to 16N genomic copies in the developmental stages assayed (Bhosale et al., 2018). Although endoreduplication is thought to increase transcription rates (Bourdon et al., 2012), general transcription might decrease as hair-cell-specific genes become more highly expressed during hair cell differentiation. Single-cell RNA-seq affords us the opportunity to explore whether transcription rates differ across development. Single-cell RNA-seq can measure both relative expression (as in bulk RNA-seq) and the total number of RNA molecules per cell. The total amount of cellular mRNA was drastically reduced across hair cell development (**Figure 4D**). This result may be due to technical bias; for example, gene expression in larger, endoreduplicated cells may be more difficult to assess with this droplet-based method. If so, the observed reduction in captured transcripts should affect all genes more or less equally. Alternatively, this observation may reflect hair cell differentiation, whereby transcription of hair-cell-specific genes should remain unaffected or increase over pseudotime. Our results support the latter scenario as transcription of hair-cell-specific genes appears to increase over pseudotime, consistent with these cells undergoing differentiation towards terminally differentiated hair cells (**Figure 4E, Supplemental Figure 7A**).

To further explore this transcriptional dynamic, we calculated RNA velocity (La Manno et al., 2018), a measure of the transcriptional rate of each gene in each cell of the hair cell cluster. RNA velocity takes advantage of errors in priming during 3’ end reverse transcription to determine the splicing rate per gene and cell. It compares nascent (unspliced) mRNA to mature (spliced) mRNA; an overall relative higher ratio of unspliced to spliced transcripts indicates that transcription is increasing. In our data, only ~4% of reads were informative for annotating splicing rates, a lower percentage than what has been used in mammalian cells for velocity analyses, and thus our results may be less reliable. Based on data for 996 genes, mean RNA velocity increased across pseudotime (**Supplemental Figure 7B,** p = 2.2 e-16 linear model, rho = 0.73). This increase in velocity was associated with the predicted changes in endoreduplication (Bhosale et al., 2018), especially between the 4N and 8N stages (**Supplemental Figure 7C,** Tukey’s multiple comparison p-value = 0.0477).

We also observed developmental signals in other cell types, including cortex and endodermis (**Figure 5A-D, Supplemental Figure 8**). Combined single-cell expression values for cortex cells highly correlated with those from bulk protoplasts sorted for expression of the *COR* cortex marker gene (**Figure 5B**, R^2^ = 0.74, rho = 0.86). As Monocle 3 did not identify a trajectory for cortex cells in the context of all cells, we isolated the cortex cells and re-performed UMAP dimensionality reduction, clustering, and graph embedding (**Supplemental Data Set 1**). Each cortex cell was assigned a developmental stage based on its Spearman’s rank correlation with bulk expression data (Brady et al., 2007; Cartwright et al., 2009). Cortex cells with the earliest developmental signal were designated as the root of the cortex trajectory, and pseudotime was assigned to the remaining cortex cells based on their distance from the root (**Figure 5A-D, Supplemental Figure 6**). As pseudotime increased for cortex cells, so did their age, indicating good agreement of the trajectory with developmental bulk RNA-seq data. Although we observed some decrease in total RNA expression and increased expression in cell-type specific genes for endodermis, we did not see a clear pattern of change in total RNA across cortex pseudotime (**Supplemental Figures 8 & 9**).

**Figure 5.**
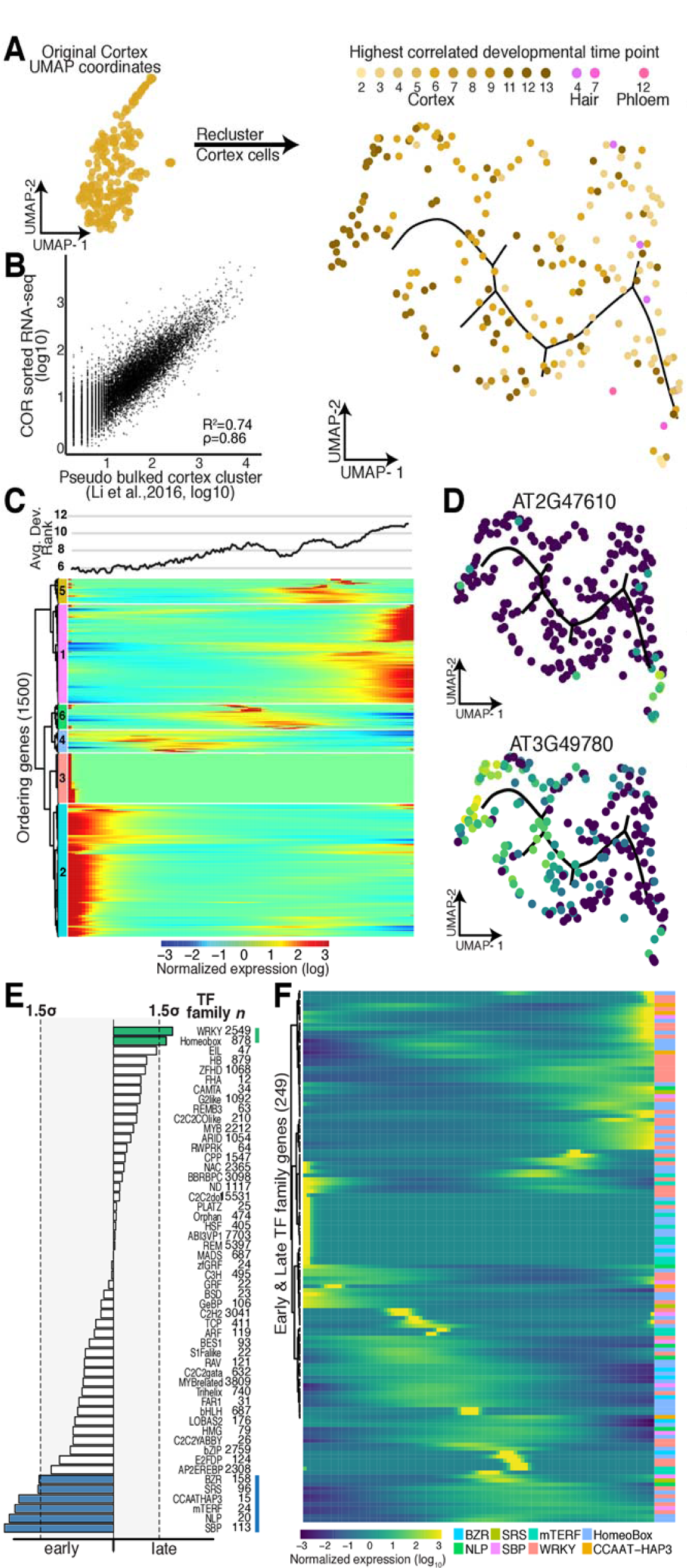
Developmental trajectory of cortex cells. **(A)** Cortex cells were re-clustered to create a trajectory, in which each cell was assigned a developmental time point and identity (shades of yellow, brown, pink) based on the highest Spearman’s rank correlation of a cell’s gene expression with prior sorted bulk data (Brady et al., 2007; Cartwright et al., 2009). **(B)** Comparison of pseudo-bulk expression dala Irom cells annotated as cortex cells with bufk expression data from protoplasts sorted for expression of the cortex marker gene COR (Li et al., 2016). **(C)** Cells were ordered in pseudotime; columns indicate cells and rows the expression of the 1500 ordering genes. Rows were grouped based on similarity in gene expression, result-in g in 6 clusters (indicated left), with genes in clusters 2 and 3 expressed early in pseudotime and genes in cluster 1 expressed late. Cortex cells with the earliest developmental signal (Brady et al., 2007; Cartwright et al, 2009) were designated as the root of the trajectory. The graph above represents the average best-correlation of developmental stage (Brady et al._t_ 2007; Cartwright et al., 2009) in a scrolling window of 20 cells with pseudotime, showing the expected increase in developmental age with increasing pseudotime. **(D)** Exampies of an early and a late expressed novel cortex-ceil-specific gene. Gene expression in each cell is superimposed onto the UMAP cluster and trajectory, with lighter colors indicating higher gene expression. **(E)** Different transcription factor motifs reside in the 500 bp upstream regions of genes expressed early (clusters 2, 3) compared to genes expressed [ate (cluster 1)· Transcription factor motifs specific to early cortex cells are denoted with blue bars, those for late cortex cells with green bars, bar length indicates motif frequency. Thresholds on either side (grey box, dotted lines) refer 10 1.5 standard deviation above mean motif frequency. (F) Expression of individual members of transcription factors families highlighted in D across pseudotime identifies candidate factors driving early or late gene expression.

We asked whether we could assign the transcription factors that drive gene expression along these developmental trajectories in early and late hair, cortex, and endodermis cells. As before, we first analyzed transcription factor motif enrichments and then explored the expression of the corresponding transcription factor gene families. Indeed, for most developmentally enriched transcription factor motifs, we could pinpoint candidate transcription factors that are expressed either early or late. For example, the *AP2/EREBP* (APETALA2/ethylene responsive element binding protein) transcription factor family is one of the largest in *A. thaliana* (Riechmann et al., 2000), with nearly 80 covered in our data set; of these, only four (AT2G25820, At5G65130, AT1G36060, AT1G44830) showed strong expression in late hair cells (**Figure 4F, G, Supplemental Figure 10**). One of these, AT1G36060 (Translucent Green), regulates expression of aquaporin genes (Zhu et al., 2014). Overexpression of this gene confers greater drought tolerance (Zhu et al., 2014), consistent with its expression in older hair cells. Similar examples of developmental stage-specific motif enrichments with corresponding transcription factor expression were also found for cortex and endodermis (**Figure 5E, F**, **Supplemental Figure 8, Supplemental Figure 10**).

### Branch points in developmental trajectories mark developmental decisions

Although a developmental trajectory that reflects the differentiation from early to late cells within a cell type should be branchless, we did observe some branch points, for example in Louvain component 8, affording us the opportunity to explore their biological relevance. As discussed, Louvain component 8 contains early non-hair cells and likely some lateral root cap cells. To further explore the cells within the branch, we performed a principal graph test, comparing their expression profiles to those of cells elsewhere in the cluster (**Figure 6A**). We found that cells within the branch were significantly enriched for expression of genes involved in cell plate formation, cytokinesis and cell cycle. We explored this enrichment for cell cycle annotations by comparing expression of previously identified core cell cycle genes (Gutierrez, 2009) in cells within the branch to cells in the rest of the cluster, finding many core cell cycle genes, in particular many G2 genes, to be specifically expressed in branch cells (**Figure 6B**). Among these genes were several of the cyclin-dependent kinase B family members that direct the G2 to M transition. Two cyclin-dependent kinase subunits (*CKS1* and *CKS2*), thought to interact with several *CDK* family members, were also specifically expressed in branch cells (Vandepoele et al., 2002). Other branch-cell-specific genes included *AUR1* and *AUR2*, both involved in lateral root formation and cell plate formation (**Figure 6C**, Van Damme et al., 2011). Louvain component 9 also showed a strong, but short branching point. We did not find any biological processes enriched in genes expressed specifically in this short branch; however, one gene whose expression is known to be affected by protoplasting was specifically expressed in these cells, perhaps reflecting that cells within this branch were more stressed by our experimental procedure (data not shown).

**Figure 6.**
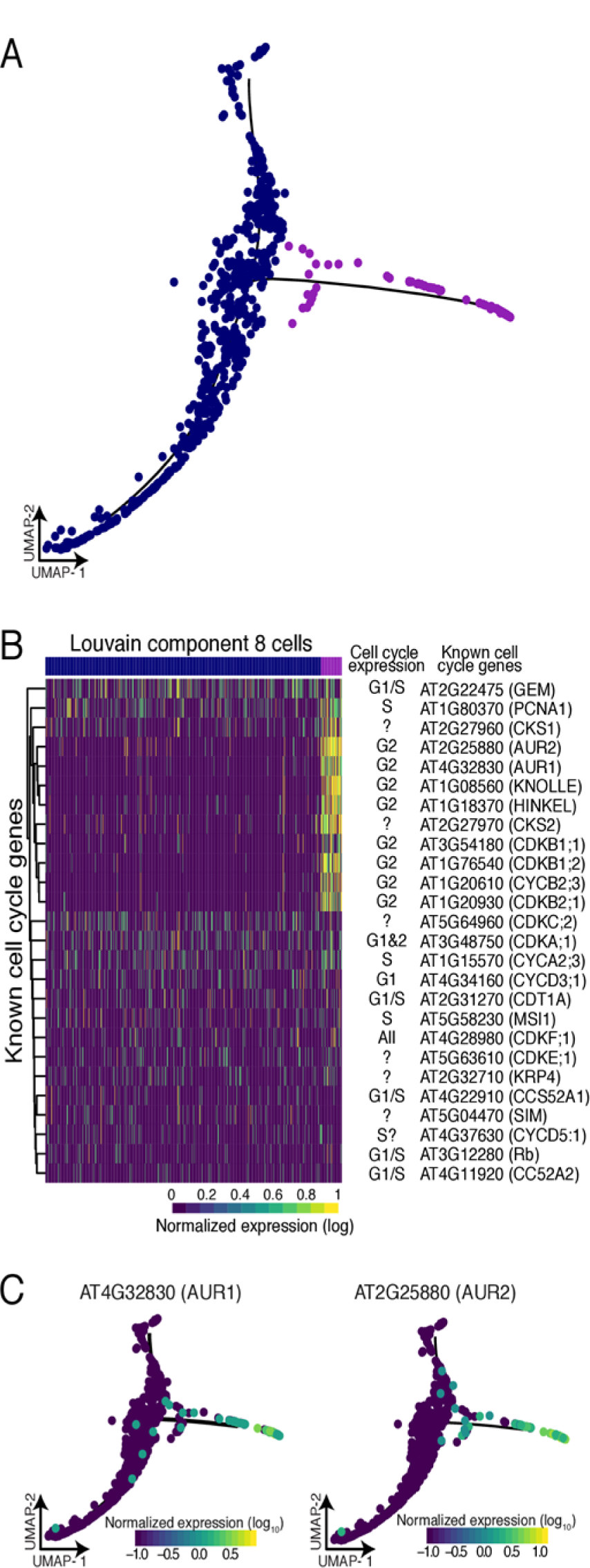
Branch analysis reveals actively dividing cells. **(A)** The 70 cells that resided in the branch of Louvain component 8 (purple) show significant branch-specific expression of genes enriched lor cell cycle function. **(B)** Comparison of all known cell cycle genes with expression in at least 5% of cells in Louvain component 8. Known cell cycle expression is denoted for each gene, if unknown ‘?’. **(C)** Two kinases, AUR1 and AUR2, were specifically expressed in branch cells. These genes are involved in cell plate formation and lateral root formation.

### Heat-shocked root cells show subtle expression differences among cell types

A major question in studying plant responses to abiotic stress, such as heat or drought, is the extent to which such responses are non-uniform across cell types. Canonically, the heat stress response is characterized by rapid and massive up-regulation of a few loci, mostly encoding heat shock proteins, with dramatic down-regulation of most other loci, in part because of altered mRNA splicing and transport (Saavedra et al., 1996; Yost and Lindquist, 1986, 1988). In plants, a set of 63 genes, most encoding heat shock proteins, show extreme chromatin accessibility at both promoter and gene body upon heat stress, consistent with their high expression (Sullivan et al., 2014). In mammals and insects, not all cells are competent to exhibit the hallmarks of the heat shock response (Dura, 1981; Morange et al., 1984); specifically, cells in early embryonic development fail to induce heat shock protein expression upon stress.

We explored whether all cells within developing roots were capable of exhibiting a typical heat shock response. To do so, we applied a standard heat stress (45 min, 38°C) to eight-day-old seedlings, harvested their roots along with roots from age- and time-matched control seedlings, and generated protoplasts for single-cell RNA-seq of both samples. For the control sample, we captured 1076 cells, assaying expression for a median 4,079 genes per cell and a total of 22, 971 genes; 82.7% of reads mapped to the TAIR10 genome assembly. The results for these control cells were similar to those described earlier with regard to captured cell types, proportion of cell types (*e.g.* 28.8% vs. 34% annotated hair cells and 9.7% vs. 7.2% endodermis cells), and correlation of gene expression (R^2^ = 0.86 for the 21,107 genes captured in both experiments). For the heat shock sample, we captured 1,009 cells, assaying expression for a median 4,384 genes per cell and a total of 21,237 genes; 79.8% of reads mapped to the TAIR10 genome assembly.

Due to global gene expression changes upon heat shock, we could not simply assign cell and tissue types as before for heat-shocked cells. The overwhelming impact of heat shock was also apparent when comparing the first and second highest cell type and developmental Spearman’s rank correlations for control cells and heat-shocked cells. Upon heat shock, many cells, especially those with non-hair, phloem and columella as their highest rank, commonly showed as their second highest rank a different cell type instead of another developmental time point of the same cell type as observed in control cells (**Supplemental Figure 11A**). Unsurprisingly, the drastic changes in gene expression led to cells being embedded in UMAP space primarily as a function of treatment, making direct comparisons of treatment effects on any one cell type impossible (**Supplemental Figure 11B**). To enable such comparisons, we used a mutual nearest neighbor to embed cells conditioned on treatment in UMAP space (Haghverdi et al., 2018). The mutual nearest neighbor method was originally developed to account for batch effects by identifying the most similar cells between each batch and applying a correction to enable proper alignment of data sets. Here, we employ this technique to overcome the lack of marker expression in our heat-shock treated cells and match them to their untreated counterpart based on overall transcriptome similarity (**Figure 7A**). This procedure yielded corresponding clusters in control and heat-shocked cells, albeit with varying cell numbers for most (**Supplemental Figure 11C, Supplemental Table 2**).

**Figure 7.**
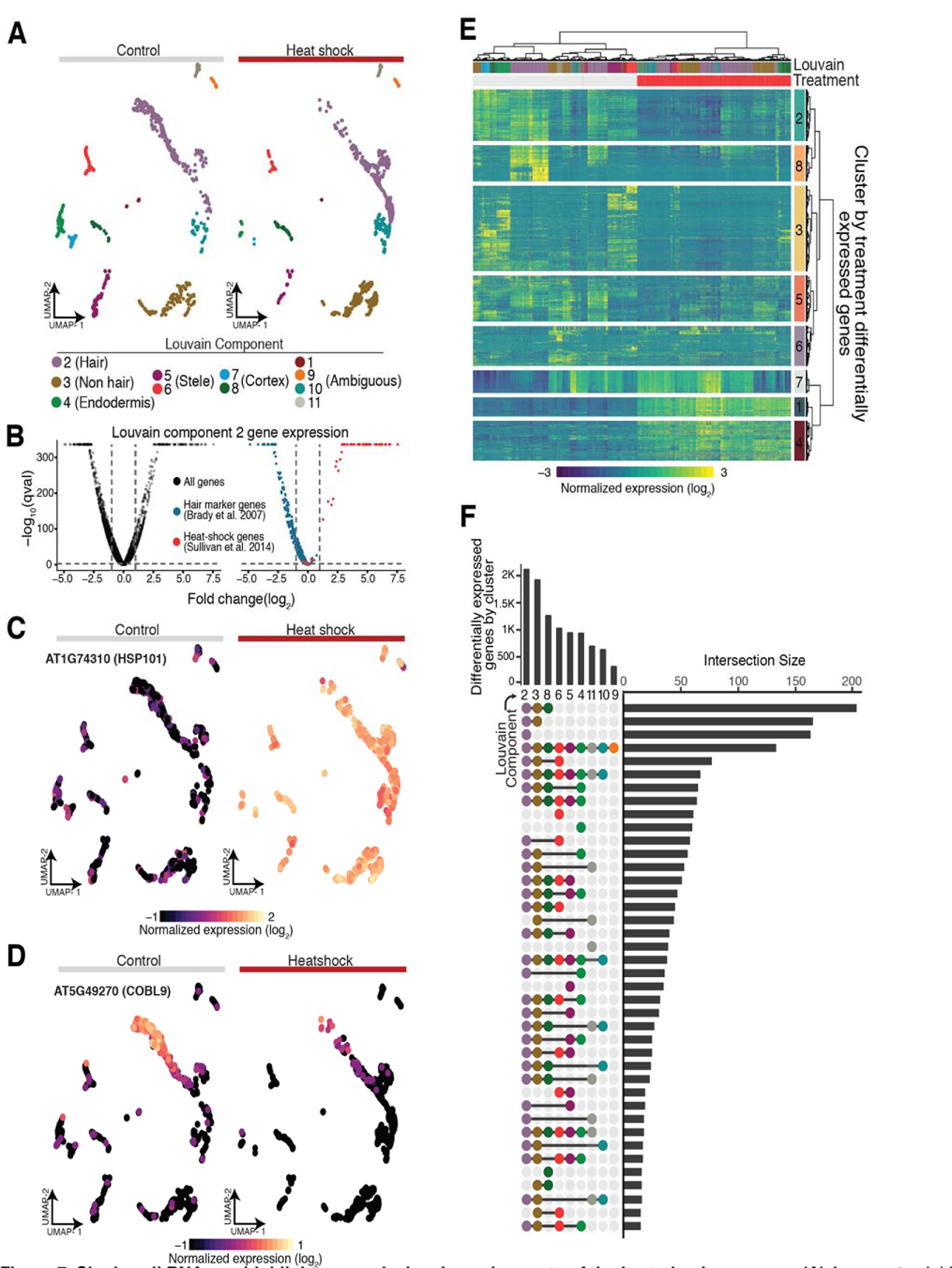
Single cell RNA-seq highlights canonical and novel aspects of the heat shock response. **(A)** A nearest neighbor approach aligns control and heat-shocked cells in a UMAP embedding to allow for concomitant cluster/cell type assignment. **(B)** Volcano plots of average gene expression change upon heat shock within Louvain component 2 for all genes (black), known hair marker genes (blue) and heat-shock signature genes (red). **(C)** HSP101, a signature heat shock gene, shows dramatic increase of expression in all cell types in all heat shock. **(D)** COBL9, a well-studied hair marker gene, is strongly repressed upon heat shock. **(E)** Heat map of of differentially expressed genes upon heat shock (top red bar; control, top gray bar), hierarchically clustered by both cells and genes (FDR < 0.1% and absolute value of the log2 fold change > 1), **(F)** Upset plot (Lex et al., 2014) of the number of differentially expressed genes as a function of heat shock for each Louvain cluster in our UMAP embedding (bars on top) along with the number of the intersect of differentially expressed genes between Louvain clusters (bars on the right). A surprising number of differentially expressed genes were specific to certain clusters (single dot in vertical row of dots).

In response to stress, organisms are thought to upregulate stress genes and to specifically downregulate genes involved in growth and development to optimize resource allocation. In response to heat stress, this presumed ‘dichotomy’ in gene expression is mirrored by the rapid localization of RNA polymerase II to the heat shock gene loci and its depletion elsewhere in the genome (Teves and Henikoff, 2011). Our data provide strong evidence of this regulatory trade-off at the level of individual cells. Using hair cells (Louvain component 2) as an example, we found that hair-cell-specific genes are overwhelmingly repressed and that heat shock genes are upregulated, often dramatically so (**Figure 7B-D)**. Indeed, *HSP101*, the most highly expressed and chromatin-accessible gene upon heat shock in previous studies (Sullivan et al., 2014), was strongly expressed across all clusters while expression of the hair marker gene *COBL9* decreased dramatically upon stress (**Figure 7C, D**).

Having established comparable clusters, we next identified genes that were differentially expressed as a function of treatment and cluster identity, excluding those with less than 15 cells in either control or heat shock conditions. This analysis identified 8,526 genes (FDR < 0.1%) whose expression was altered by heat shock treatment in one or more clusters; of these, 2,627 genes were up- or downregulated at least 2-fold (**Figure 7E, Supplementary Data Set 5**, FDR < 0.1% and absolute value of log2 fold change > 1). As for hair cells (**Figure 7B**), cell-type marker genes for all clusters were enriched among the downregulated genes upon heat shock. To identify cluster-specific differences in the response to heat shock, we compared gene expression of cells within individual clusters to the rest of the cells across treatments. We observed the largest number of cluster-specific gene expression changes in hair, non-hair and cortex cells (**Figure 7F**). As these cell types are the three outermost cell layers of the root, they may be exposed more directly to the heat shock and respond more quickly. Genes differentially expressed in hair cells (Louvain component 2) upon heat shock were enriched for ribosome associated genes and RNA methylation. Stele cells (Louvain component 6) showed differential expression of genes involved in cell wall organization and biogenesis, and endodermis cells (Louvain component 4) showed differential expression of genes involved in response to external, chemical and stress stimuli as well as nitrate and anion transport (**Figure 7F**).

The expression of heat shock proteins protects cells from heat shock and aids their recovery (Parsell et al., 1993; Parsell and Lindquist, 1993; Queitsch et al., 2000). We were interested in whether we could detect cluster- and cell-type-specific differences in the canonical heat shock response. In principle, such differences could be exploited to alter heat shock protein expression in a cell-type-specific manner to boost plant heat and drought tolerance without pleiotropically decreasing whole-organism fitness. To address such possible differences, we focused on genes that from bulk analyses have differential expression upon heat shock (1783 genes) or reside near regulatory regions that change in accessibility upon heat shock (1730 genes) (Alexandre et al., 2018; Sullivan et al., 2014). Although these gene sets overlap (942 genes), they contain complementary information, as changes in accessibility do not necessarily translate into altered expression, and vice versa (Alexandre et al., 2018). In our single-cell expression analysis, we identified 752 of 1783 heat-responsive genes as differentially expressed upon heat shock, and 564 of 1730 genes near dynamic regulatory regions as differentially expressed. We hierarchically clustered control and heat shock-treated single-cell transcriptomes for both gene sets (**Supplemental Figure 12A, C**), resulting in several gene clusters with distinct expression patterns. Overall, cellular responses were dominated by the canonical heat-shock response, as visualized in cluster 4 (**Supplemental Figure 12A**) and cluster 2 (**Supplemental Figure 12C**). The 63 genes showing extreme accessibility and high expression upon heat shock (Sullivan et al., 2014) are largely contained in these two clusters (**Supplemental Figure 12A**, cluster 4, 49 of 63; **Supplemental Figure 12C**, cluster 2, 42 of 63).

Our analysis also revealed subtle but significant differences among some tissue types (**Supplemental Figure 12A, B,** *e.g.* clusters 3 and 8, **Supplemental Figure 12C, D,** *e.g.* clusters 5 and 7, **Supplemental Data Set 6**). Although most of these gene clusters were not enriched for specific annotations, cluster 8 genes were associated with rRNA metabolic processes (p- value = 0.048) and cluster 5 genes (**Supplemental Figure 12A, B**) were enriched for transport genes (p-value = 0.045). These results demonstrate both the promise and the challenges inherent in comparing single-cell data across different conditions and treatments.

## DISCUSSION

Here, we use *A. thaliana* roots to establish both experimental and analytic procedures for single-cell RNA-seq in plants. Using Monocle 3, we could assign over 3000 cells to expected cell and tissue types with high confidence. In particular, cortex, endodermis and hair cells were easily identified. However, distinguishing other cell types was challenging. For example, non-hair and columella cells had high similarity in their expression profiles, consistent with their correlation in bulk expression data (Brady et al., 2007; Cartwright et al., 2009). Similarly, it was difficult to designate cells in Louvain component 8 as early non-hair cells, as these cells showed overlapping expression signatures for early non-hair cells, lateral root caps, and epidermis cells before differentiation to hair and non-hair cells. These Louvain component 8 cells were difficult to distinguish further with the sparse expression data typical for single cell analysis, however we postulate that in fact the branch of component 8 may actually be the root of the trajectory and are cells dividing out of the epidermis/root cap precursor and cells either become root cap cells or epidermis.

We also could not initially split stele tissue into individual cell types, likely because the difficulty of digesting the cell walls of the tightly packed vascular bundle resulted in fewer cells than expected (Brady et al., 2007; Cartwright et al., 2009). However, analyzing stele cells separately yielded 6 sub-clusters, which correspond to known vasculature cell types. Our approach to annotate these sub-clusters exemplifies the ad hoc nature of current single-cell genomics studies, which require all available sources of information to be exploited to interpret the genomic data. Neither Spearman rank correlations with sorted bulk RNA-seq data nor microarray expression data yielded obvious cluster identities. However, mean expression values of genes known to be expressed in vasculature cell types allowed us to assign the stele sub-clusters.

We identified hundreds of novel genes with cell-type-specific and tissue-type-specific expression, which may allow the generation of new marker lines for detailed genetic analyses. These genes, together with cluster-specific enriched transcription factor motifs and their corresponding transcription factors, are candidates for driving differentiation and cell-type identity. Similarly, the developmental trajectories we identified highlight the potential of single cell transcriptomics to advance a high resolution view of plant development. These trajectories can be detected without the use of spatial information because plants have a continuous body plan with new cells continuously arising while older cells persist. Additionally, while this study allowed us to infer transcription factor motifs and candidate transcription factors, future analyses with greater numbers of cells than assayed here may include combinatorial expression of multiple transcription factor family members.

We explored the relationships of endoreduplication, transcriptional rates, and differentiation to find that transcriptional rates, measured as mRNA velocity, increase with increasing ploidy. However, this transcriptional increase appears to be limited to genes specifically expressed in hair cells, as overall levels of RNA decreased over pseudotime. These observations are consistent with hair cells becoming more specialized and moving towards a terminally differentiated state over time. However, this phenomenon of increasing specialization was not as apparent in other cell types. This difference may be due to biological causes, such as the higher rates of endoreduplication in hair cells, or to technical causes, such as the better clustering and trajectory of hair cells compared to the other cell types assayed.

By allowing trajectories with side branches, we discovered that branch points can mark developmental decisions. In Louvain component 8, the small but distinct cell-cycle enriched branch may mark lateral root primordia cells differentiating into epidermal cells or epidermal/lateral root precursor cells. Cells within this branch express many cell cycle genes, among them members of the *CDKB* family that govern the G2 to M transition. Moreover, these cells specifically express the *AUR1* and *AUR2* genes, which function in cell plate formation; plants with mutations in these genes lack lateral roots (Van Damme et al., 2011). Although expression of cell cycle genes may persist in non-dividing cells because of their roles in endoreduplication, *AUR1* and *AUR2* expression (and cell plate formation) should not persist, consistent with our speculation that the cells within this branch are actively dividing cells in the G2 to M transition (Gutierrez, 2009).

We explored the *A. thaliana* heat shock response with single-cell RNA-seq because not all cells and tissues are equally competent to respond to stress. By identifying plant cell types that most strongly respond to abiotic stresses such as heat, drought, and nutrient starvation, ultimately we may be able to genetically manipulate relevant cell types to generate stress-tolerant crops without pleiotropically affecting plant fitness and yield. Although all heat-shocked cells showed gene expression changes typical of the canonical heat shock genes, we detected subtle but highly significant expression differences among cells and tissue types for other genes. Thus, single-cell transcriptomics across stress conditions holds potential for future crop breeding and genetic engineering. However, such analyses require much larger numbers of cells than currently accessible by droplet-based methods. Moreover, such analyses should focus on treatments that are less overwhelmed by a strong canonical signal to increase resolution in detecting cell-type-specific differences.

In this study, we relied on the extensive and detailed expression data for bulk *A. thaliana* cell and tissue types to establish the validity of our approaches. The overwhelming correspondence of our findings with these and other data derived from traditional molecular genetics provides confidence that less well-characterized *A. thaliana* tissues and other plants, including crops, will be amenable to these approaches. Thus, continued progress on single-cell RNA-seq experiments should have a major impact on the analysis of plant development and environmental response.

## METHODS

### Plant Material and Growth Conditions

*Arabidopsis thaliana* Col-0 seedlings were grown vertically at 22°C, on 1xMS + 1% sucrose plates covered with one layer of filter paper. Seven or eight days-old seedlings (LD, 16h light/8h dark, ~100 μmol m2 s) were collected around ZT3, and the roots/shoots excised with a sharp razor blade. For the heat-shock, seedling plates were transferred from 22°C to 38°C for 45 min (Conviron TC-26, light ~100 μmol m2 s), and the roots harvested immediately after.

### Protoplast Isolation

Protoplast isolation was done as previously described (Bargmann and Birnbaum, 2010), with slight modifications. Briefly, 1 g of whole-roots was incubated in 10 ml of protoplasting solution for 1.5 h at 75 rpm. After passing through a 40 μm strainer, protoplasts were centrifuged at 500 g for 5 min and washed once in protoplasting solution without enzymes. Final suspension volume was adjusted to a density of 500–1,000 cells/μl. Protoplasts were placed on ice until further processing.

### Single-cell RNA-seq protocol

Single-cell RNA-seq was performed on fresh Arabidopsis root protoplast using the10X scRNA-seq platform, the Chromium Single Cell Gene Expression Solution (10X Genomics).

### Data Analysis

#### Estimating gene expression in individual cells

Single-cell RNA-seq reads were sequenced and then mapped to the TAIR10 Arabidopsis genome using Cellranger (version 2.1.0) (https://support.10xgenomics.com/single-cell-gene-expression/software/pipelines/latest/what-is-cell-ranger). Cellranger produces a matrix of UMI counts where each row is a gene and each column represents a cell. The ARAPORT gene annotation was used. For the heat shock analysis, reads from a control sample and reads from heat-shocked sample were aggregated using “cellranger aggr” to normalize libraries to equivalent number of mean reads per cell across libraries.

#### Running Monocle 3: Dimensionality Reduction, and Cell Clustering

The output of the cellranger pipeline was parsed into R (version 3.5.0) using the cellranger R kit (version 2.0.0) and converted into a CellDataSet (cds) for further analysis using Monocle 3 alpha (version 2.99.1) (http://cole-trapnell-lab.github.io/monocle-release/monocle3/). All Monocle 3 analysis was performed on a High Performance Computing cluster using 128GB of RAM spread across 8 cores. The lower detection limit for the cds was set at 0.5, and the expression family used set to negbinomial.size().

We visualized cell clusters and trajectories using the standard Monocle workflow. Monocle internally handles all normalization needed for dimensionality reduction, visualization, and differential expression via “size factors” that control for variability in library construction efficiency across cells. After estimating the library size factors for each cell (via estimateSizeFactors), and estimating the dispersion in expression for each gene (via estimateDispersions) in the dataset, the top 1500 genes in terms of dispersion, *i.e*. 1500 genes with the most expression variability in our dataset, were selected to order the cells into clusters. The expression values of these 1500 genes for each cell were log-transformed and projected onto the first 25 principal components via Monocle’s data pre-processing function (preprocessCDS). Then, these lower-dimensional coordinates were used to initialize a nonlinear manifold learning algorithm implemented in Monocle 3 called Uniform Manifold Approximation and Projection, or UMAP (via reduceDimension) (McInnes and Healy, 2018). This allows us to visualize the data unto two or three dimensions. Specifically, we projected onto 2 components using the cosine distance metric, setting the parameters “n_neighbors” to 50, and “min_dist” to 0.1.

The Louvain method was used to detect cell clusters in our two dimensional representation of the dataset (partitionCells); this resulted in 11 cell clusters, or Louvain components. Cells were then clustered into “super” groups using a method derived from “approximate graph abstraction” (Wolf et al., 2018) and for each super group, a cell trajectory was drawn atop the projection using Monocle’s reversed graph embedding algorithm, which is derived from “SimplePPT” (learnGraph) (Mao et al., 2015). This yielded 6 cell trajectories.

To further analyze the clusters we annotated as stele, Clusters 3, 4, and 7 were reclustered together and were reanalyzed using Monocle 3 as previously described except the parameter “min_dist” was changed to 0.05 when the reduceDimension function was called. This revealed 6 additional sub clusters.

To further analyze the cluster we annotated as cortex, Cluster 10 was reclustered and reanalyzed using Monocle 3 as previously described except the parameters “n_neighbors” was reduced to 25. This did not reveal any sub clusters, but a trajectory was generated.

#### Estimating doublets

**S**ingle-**C**ell **R**emover of Do**ublet**s (Scrublet) was used to predict doublets in our scRNA-seq data (Available at: https://github.com/AllonKleinLab/scrublet). Using Python 3.5, Scrublet was ran using default settings as described by the example tutorial which is available as a python notebook (Available at: https://github.com/AllonKleinLab/scrublet/blob/master/examples/scrublet_basics.ipynb). The only significant change was that expected double rate was set to 0.1, in the tutorial it is 0.06.

#### Identifying Cell Types

In order to categorize the cells into cell types and to apply developmental information, a deconvolved root expression map was downloaded from AREX LITE: The <u>Ar</u>abidopsis Gene <u>Ex</u>pression Database (http://www.arexdb.org/data/decondatamatrix.zip). Using this data matrix, the Spearman’s rank correlation was calculated between each cell in our dataset and each cell type and longitudinal annotation in the data matrix (3121 × 128 Spearman’s rank correlations total). Specifically, we looked at the correlation of 1229 highly variable genes in our dataset. These 1229 genes represents the overlap between our 1500 highly variable genes and genes in the root expression map data matrix. Cells in our dataset were assigned a cell type and a developmental label based on the annotation with which each cell had the highest correlation. (*i.e.* if a cell correlated highest with endodermis cells in longitudinal zone 11, then it would be called as endodermis_11).

In addition to using the Spearman’s rank correlation to assign cells their cell type, a set of known marker genes derived from GFP marker lines of the Arabidopsis root were used to identify cell types based on the high gene expression of these marker genes. These genes were obtained from (Brady et al., 2007; Cartwright et al., 2009). Specifically Supplemental Table 2 (Cartwright et al., 2009) was used. For the analysis comparing bulk RNA and pseudo bulk scRNA-seq data, the bulk data was obtained from Li et al. 2016 (Li et al., 2016); specifically, we used Table S5 from this study. Isoforms of each gene were averaged in order to be comparable to the pseudo bulk data. Lastly, using this same bulk RNA-seq data, the Pearson correlation was calculated between each cell in our dataset and each GFP marker line. Cells in our dataset were assigned to a GFP marker line based on the GFP marker line with which each cell had the highest correlation.

#### Running Monocle 3: Identifying High Specificity Genes

In order to identify differentially expressed genes between cell clusters the Moran’s I test was performed on our UMAP (principalGraphTest), with the projection being broken up into 25 × 25 spatial units. Then marker genes were identified for each cluster, and each annotated grouping of clusters using a Moran’s I threshold of 0.1 and a qval threshold of 0.05. In order for a gene to be considered highly specific, it must have had a specificity rating of above 0.7.

#### Transcription factor motif analysis

Highly specific genes were identified for each cell cluster, and their promoters were analyzed for presence of transcription factor motifs. Promoters were defined as 500 base pairs upstream of the start site of each gene. Instances of each motif were identified using (Grant et al., 2011) at a p-value cutoff of 1e-5 for each match. The input position weight matrices for each motif were enumerated in a previous study of binding preferences for nearly all Arabidopsis transcription factors (O’Malley et al., 2016). Motif frequencies in genes specific to each cell cluster were compared to a background set of motif frequencies across all promoters in the Arabidopsis genome to determine a log2 enrichment score. TF family genes were pulled from the gene family page of TAIR10 (https://www.arabidopsis.org/browse/genefamily/index.jsp).

#### Running Monocle 3: Assigning Pseudotime

Pseudotime analysis requires the selection of a cell as an “origin” for the pseudotime trajectory. Origin assignment was based on the Spearman’s rank assignments for each cell. The following cells were used as origins for their respective cell type trajectories: cortex_2, hair_2, endodermis_2, nonHair_3. The get_correct_root_state() function was used to assign the root of a trajectory, and the orderCells() function was used to assign cells a pseudotime value.

#### Calculating total mRNA

After pseudotime analysis was performed on a cell cluster, cells were binned together such that each bin contained a similar number of cells and each bin represented cells from similar pseudotimes. The median total mRNA and the standard deviation of the total mRNA of each bin was then calculated.

#### Calculating significance with the Permutation Test

The permutation test was used to calculate the significance of the observed trends that the total mRNA of hair marker genes and hair specific genes increased as pseudotime increased in hair cells. To do this, 10000 random samplings of 441 genes (the number of hair marker genes), and 201 genes (the number of hair specific genes) were taken respectively. Next, the median total mRNA was calculated across pseudotime for each random sampling and the slope of this data was calculated using a generalized linear model. The observed slope of the marker genes and the hair specific genes was compared to the distribution of slopes generated by 10000 random samplings. No random sampling of genes had a slope that was higher than the observed slopes generated by the hair marker genes or the hair specific genes. The significance, or the p-value, of the trend seen in the hair marker genes and the hair specific genes can then be calculated simply as the proportion of sampled permutations that have a slope that is equal to or greater than slope generated by our genes of interest. This gives us a p-value of 1/10001 or roughly 1 × 10^−4^.

#### Analyzing Expression Differences Between Branches of Louvain Component 8 (Early Non-Hair)

To identify genes responsible for the branching in the pseudotime trajectory of Louvain component 8 (early non-hair), the principal graph test was used to identify genes with expression specific to the side branch vs. the main branch. Genes were considered specific if it had a specificity value above 0.8. Genes were removed from the analysis if they did not have expression in at least 10% of the cells considered and a mean expression greater than 0.25.

#### Calculating RNA velocity

We used the Velocyto R and Python packages (version 0.6 and 0.17, respectively) to estimate RNA velocity for root hair cells (La Manno et al., 2018). Matrices of spliced and unspliced RNA counts were generated from Cellranger outputs using velocyto.py CLI and “run10x” defaults. We followed the velocyto.py and velocyto.R manuals (http://velocyto.org/) and used spliced (emat) and unspliced (nmat) matrices to estimate RNA velocity. With predefined cell type annotations, we performed gene filtering with the parameter “min.max.cluster.average” set to 0.2 and 0.05 for “emat” and “nmat”, respectively. RNA velocity using the selected 996 genes was estimated with the defaults to the function gene.relative.velocity.estimates() except parameters “kCells” and “fit.quantile” which were set to 5 and 0.05, respectively. Velocity measurements for each cell were calculated as the difference between “$projected” and “$current” (with $deltaT = 1) results from the estimated velocity output.

#### Analysis of heat shock data

For each pair of cell types and for each gene cluster, we used a generalized linear model to determine the significance of an interaction between the effects of cell type and heat treatment on the normalized expression level of genes in that cluster. Then, to identify differentially expressed genes specific for every Louvain cluster we subsetted cells from every cluster that contained 15 or more cells in both control and treated conditions, estimated dispersions for each subset and tested for differential gene expression identified using the differentialGeneTest function in Monocle specifying a full model of Treatment cluster and a residual model of 1. FDR values per gene were then obtained across all tests using the Bejamini-Hochberg method. The overlap of differentially expressed genes as a function of heatshock treatment between clusters was visualized using an UpsetR plot. Briefly, a binary matrix of differentially expressed genes by cluster was generated were gene-cluster combinations were set to 1 (significant) or 0 (not significant). This matrix was then passed to the upset function from the UpsetR R package specifying 9 sets and ordering by frequency. To identify whether clusters contained subtle differences in the expression of previously identified heat shock responsive genes we tested for differential gene expression across all cells and clusters and identified the intersect between differentially expressed genes obtained from single cell profiles and previously identified dynamic changes in DHS linked genes and bulk differentially expressed genes upon heat shock. Differentially expressed genes as a function of heat-shock treatment for all cells in unison where identified using the differentialGeneTest function in Monocle specifying a full model of Treatment*UMAP cluster and a residual model of UMAP cluster. Hierarchical clustering of these DHS linked and bulk differentially expressed gene sets across control and heat-shock treated cells was performed using the pheatmap function in the pheatmap R package (version 1.0.10) specifying ward.D2 as the clustering method. Genes with similar dynamics across treatment and cell types were recovered using the cutree function from the stats package in R specifying k = 8 for both DHS linked genes and bulk differentially expressed genes. To generate signatures from these 8 groups of clustered genes we log normalized expression values using a pseudocount of 1 and for each cell calculated the mean normalized expression value across genes that belong to one of the 8 gene cluster.

## Data Availability

All sequencing data can be found on GEO at: https://www.ncbi.nlm.nih.gov/geo/query/acc.cgi?acc=GSE121619

## Supplemental Data

**Supplemental Figure 1:** General tissue and data features

**Supplemental Figure 2:** Pearson correlation to sorted RNA-seq samples

**Supplemental Figure 3:** Marker gene expression in cell type clusters

**Supplemental Figure 4:** Examples of tissue-specific gene expression

**Supplemental Figure 5:** Transcription factor family expression patterns

**Supplemental Figure 6:** Spearman’s rank correlation for each cell’s development and tissue-type

**Supplemental Figure 7:** Changes in transcription across hair development

**Supplemental Figure 8:** Developmental trajectory of endodermal cells

**Supplemental Figure 9:** Total RNA in cortex across pseudotime

**Supplemental Figure 10:** Developmental expression of individual transcription factors

**Supplemental Figure 11:** Heat-shock clustering and expression profiling

**Supplemental Figure 12:** Conditional expression in genes with dynamic chromatin accessibility during heat-shock

**Supplemental Table 1:** Bulk RNA-seq comparisons to single cell RNA-seq

**Supplemental Table 2:** Number of cells in the control vs. heatshock analysis

**Supplemental Data Set 1:** List of Ordering/ High Dispersion Genes

**Supplemental Data Set 2:** Correlation with Bulk Expression Data

**Supplemental Data Set 3:** Marker Genes

**Supplemental Data Set 4:** Novel High Specificity Genes

**Supplemental Data Set 5:** Cluster Specific Heat shock Differentially Expressed Genes

**Supplemental Data Set 6:** Generalized Linear Model pairwise test of significance between cortex, hair, and non-hair cells

## ACKNOWLEDGEMENTS

We thank members of the Trapnell Lab for input and discussion. This work was supported by NSF MCB-1516701 to CQ and NSF RESEARCH-PGR 1748843 to CQ and SF. KJB was upported by the University of Washington NIH Big Data for Genomics and Neuroscience Training Grant (T32LM012419). NIH grant U54DK107979 to CT; NIH grant DP2HD088158, RC2DK114777 and R01HL118342 to CT; and the Paul G. Allen Frontiers Group to CT.

