## Supplemental Figures 1-12, Supplemental Table 1 and 2 for "Dynamics of gene expression in single root cells of *A. thaliana*"

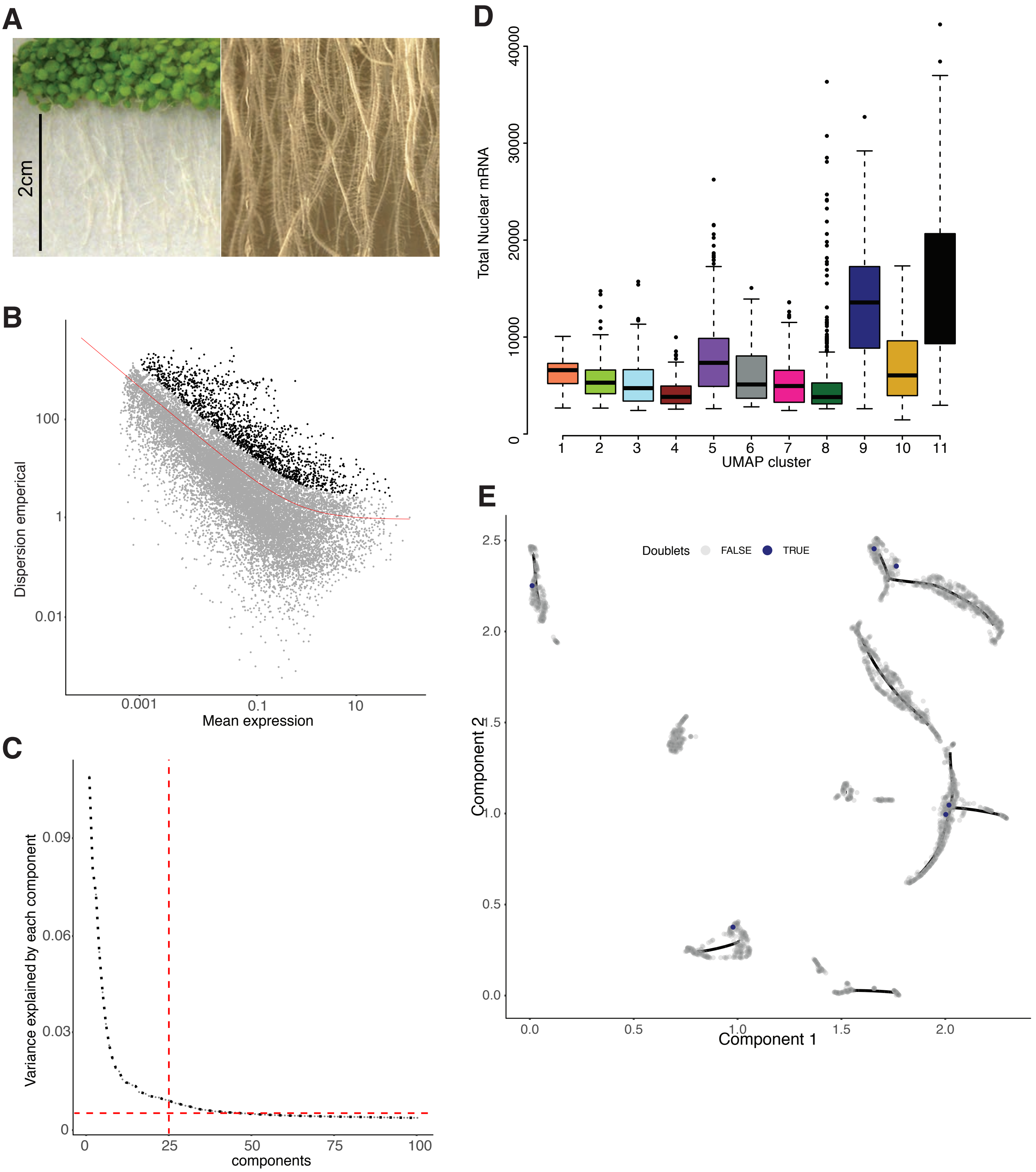


Supplemental figure 1

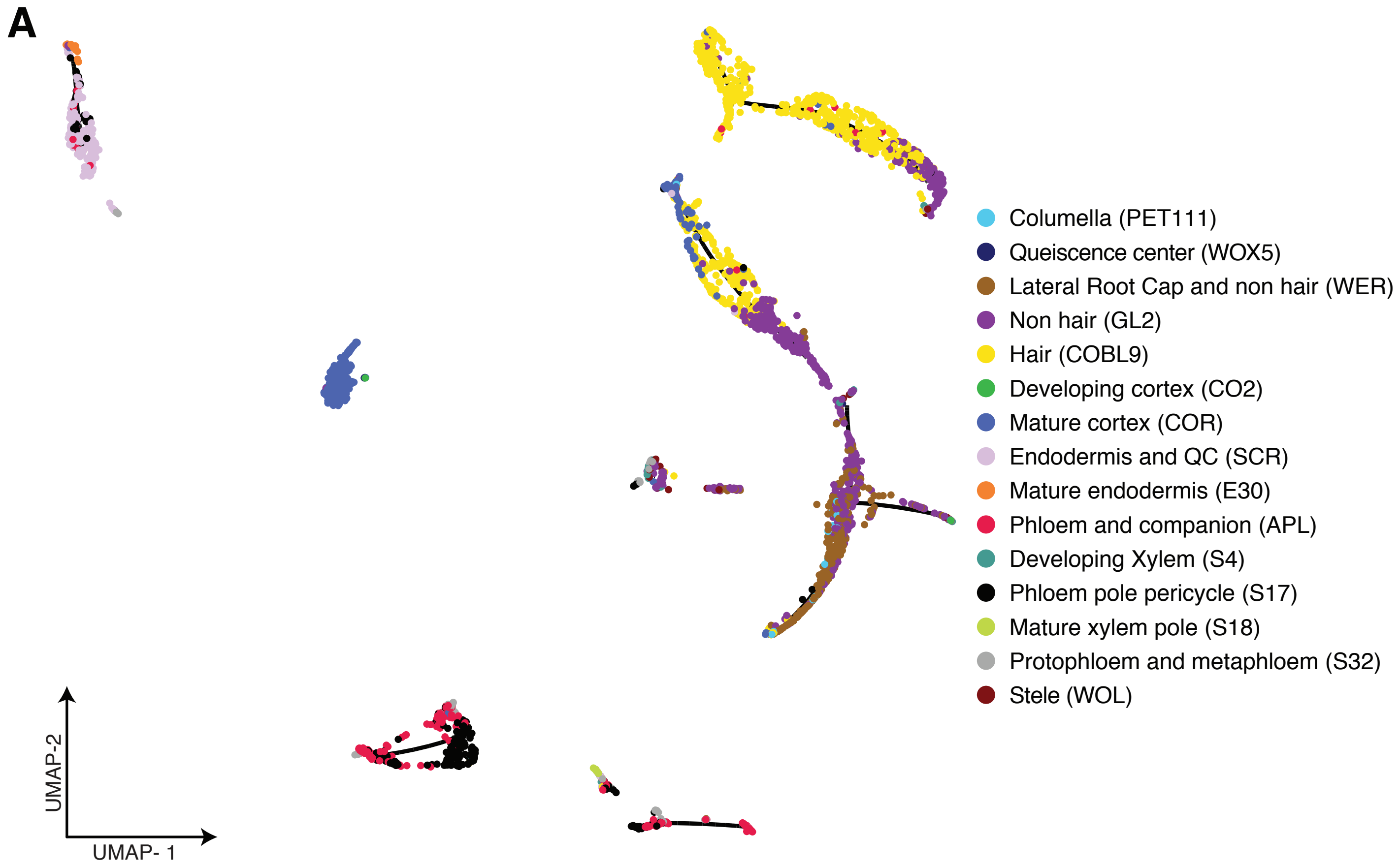


Supplemental Figure 2

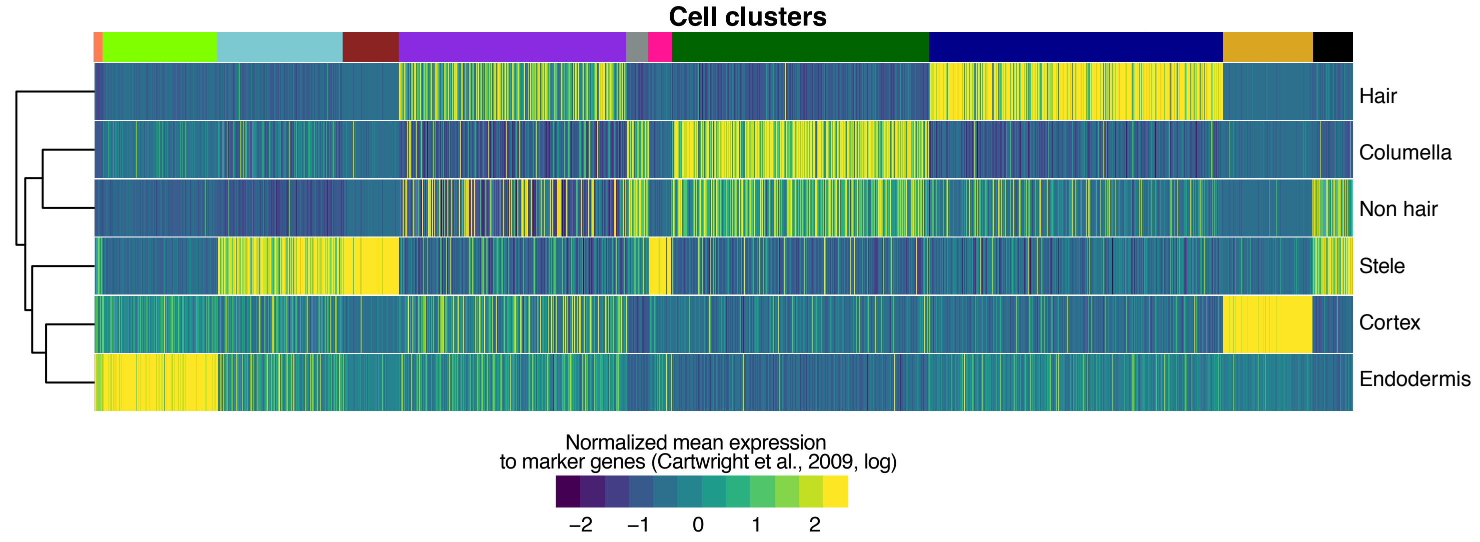


Supplemental figure 3

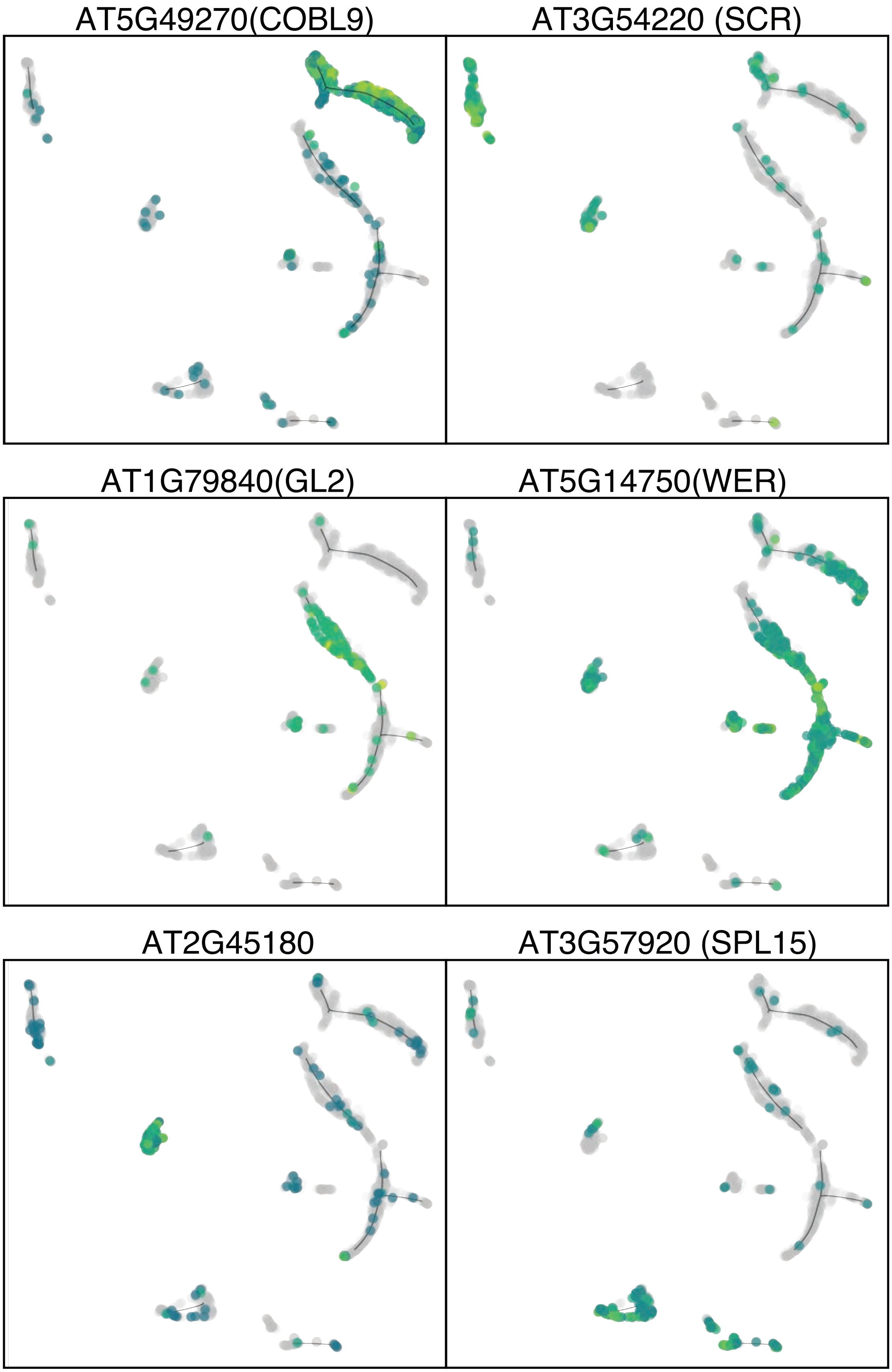


Supplemental figure 4

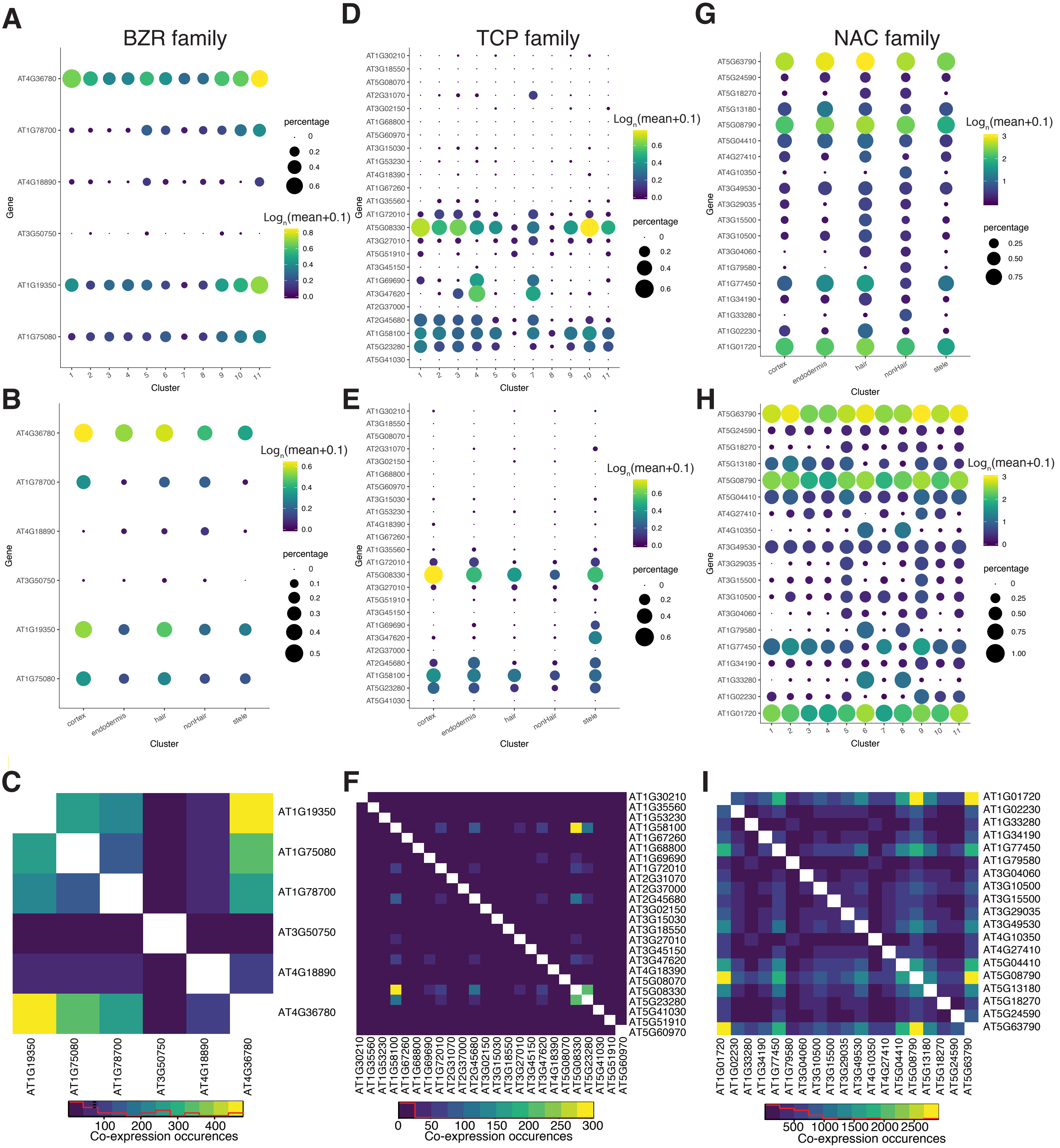


Supplemental Figure 5

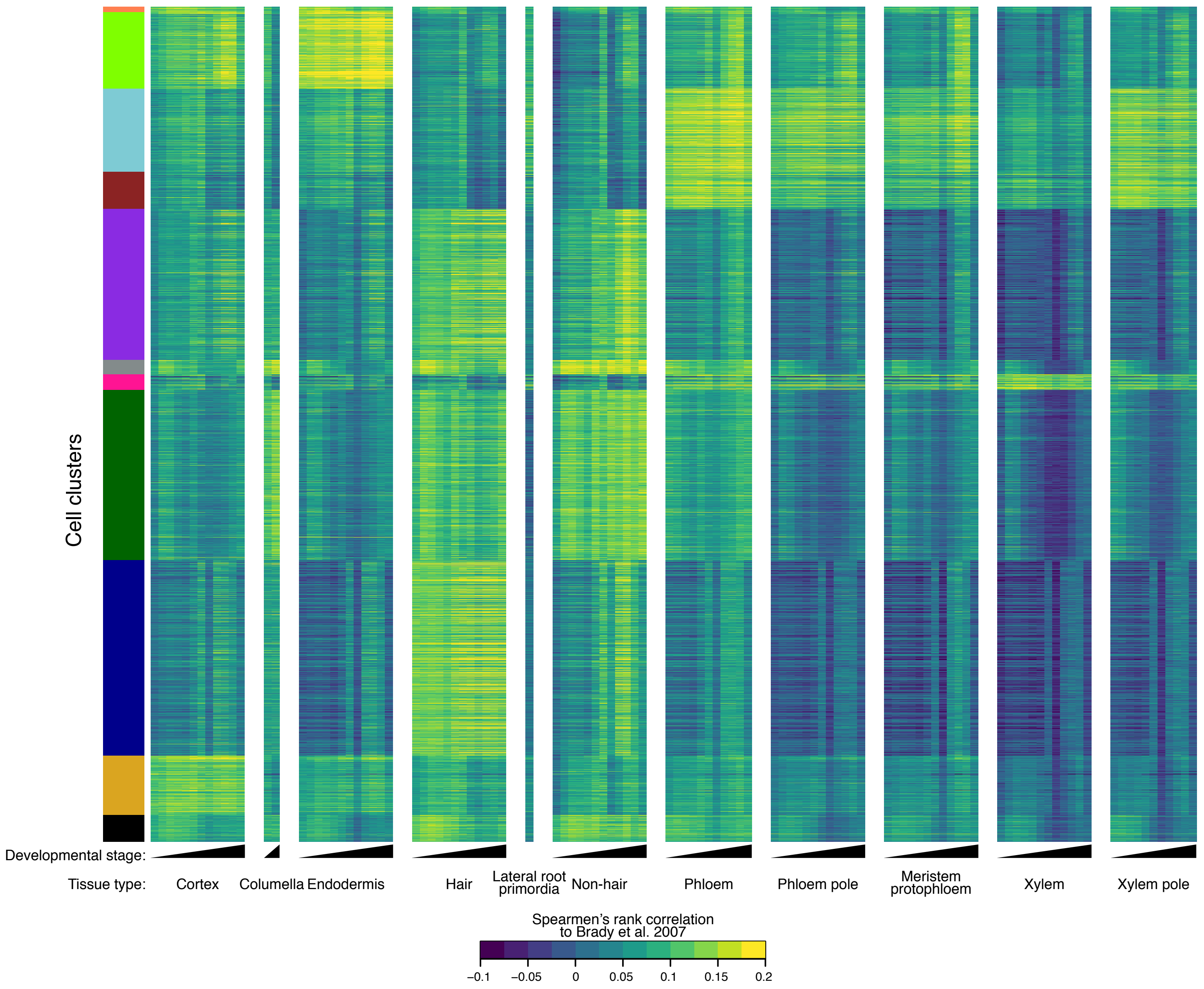


Supplemental Figure 6

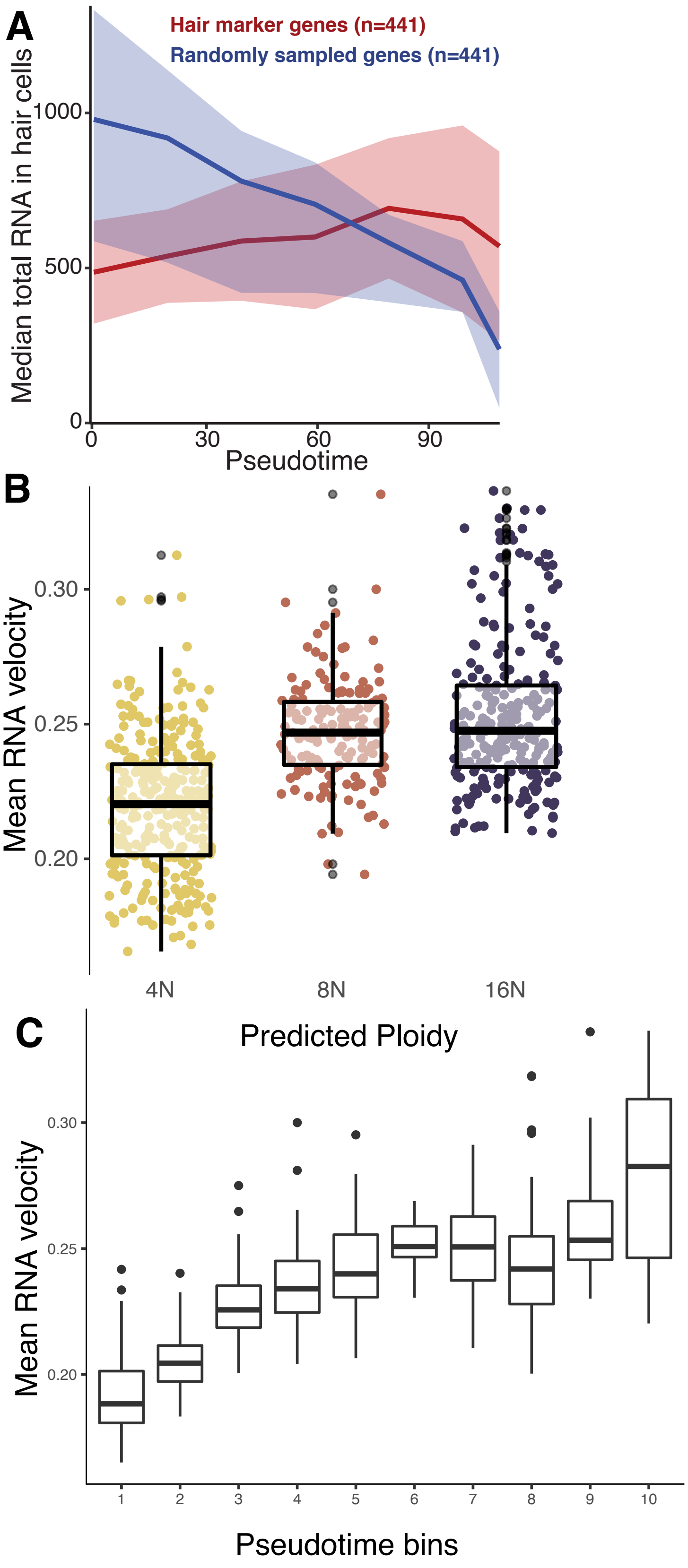


Supplemental Figure 7

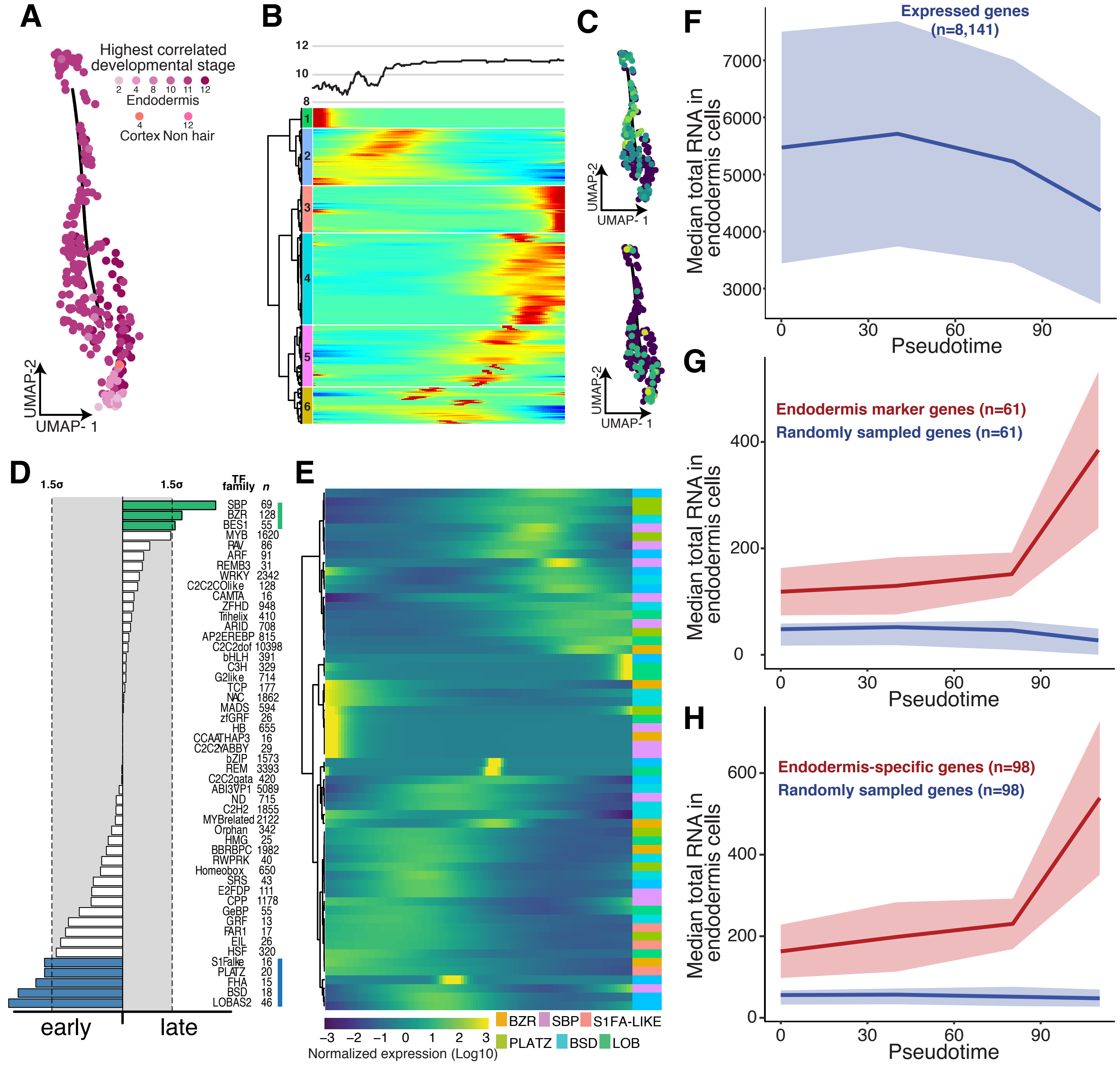


Supplemental Figure 8

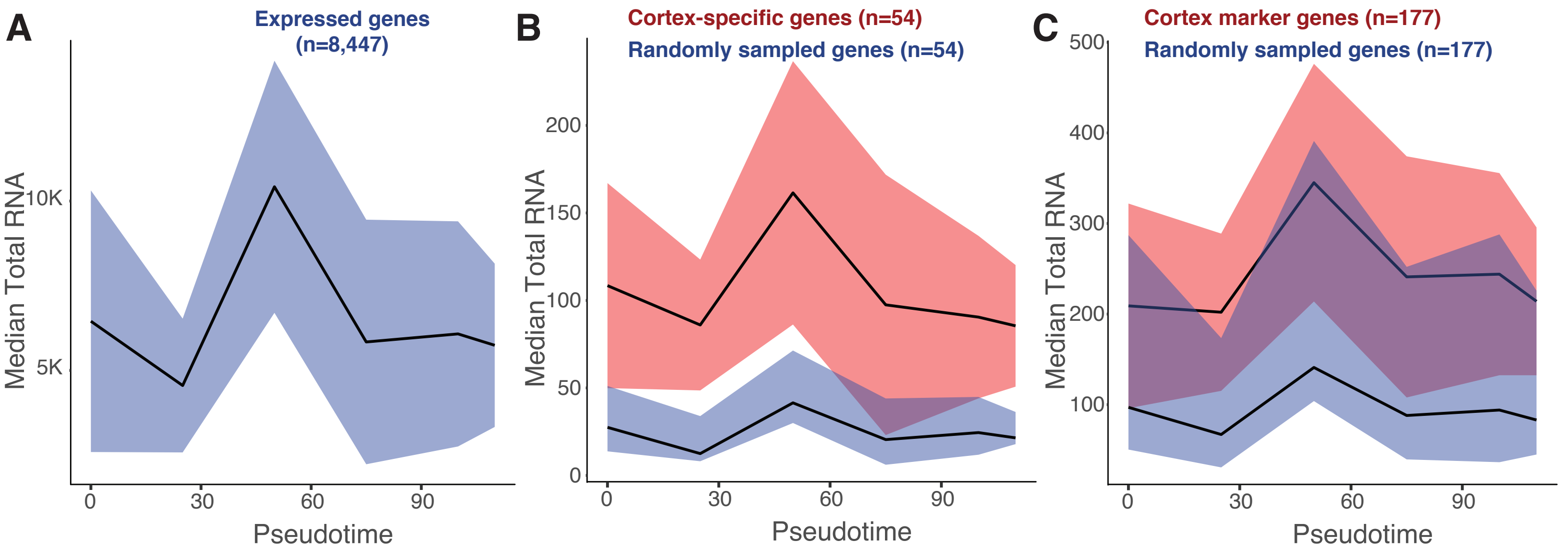


Supplemental Figure 9

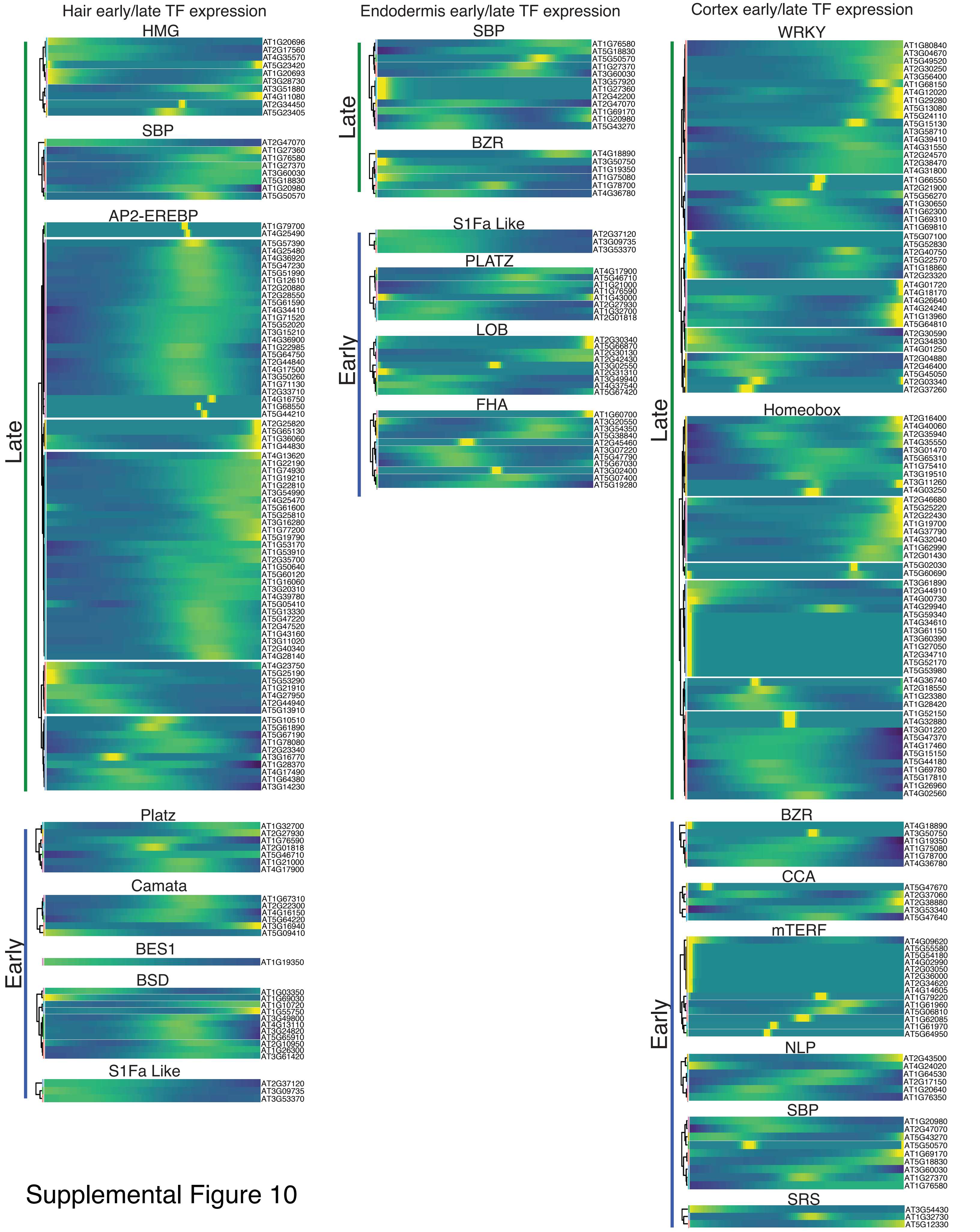


Supplemental Figure 10

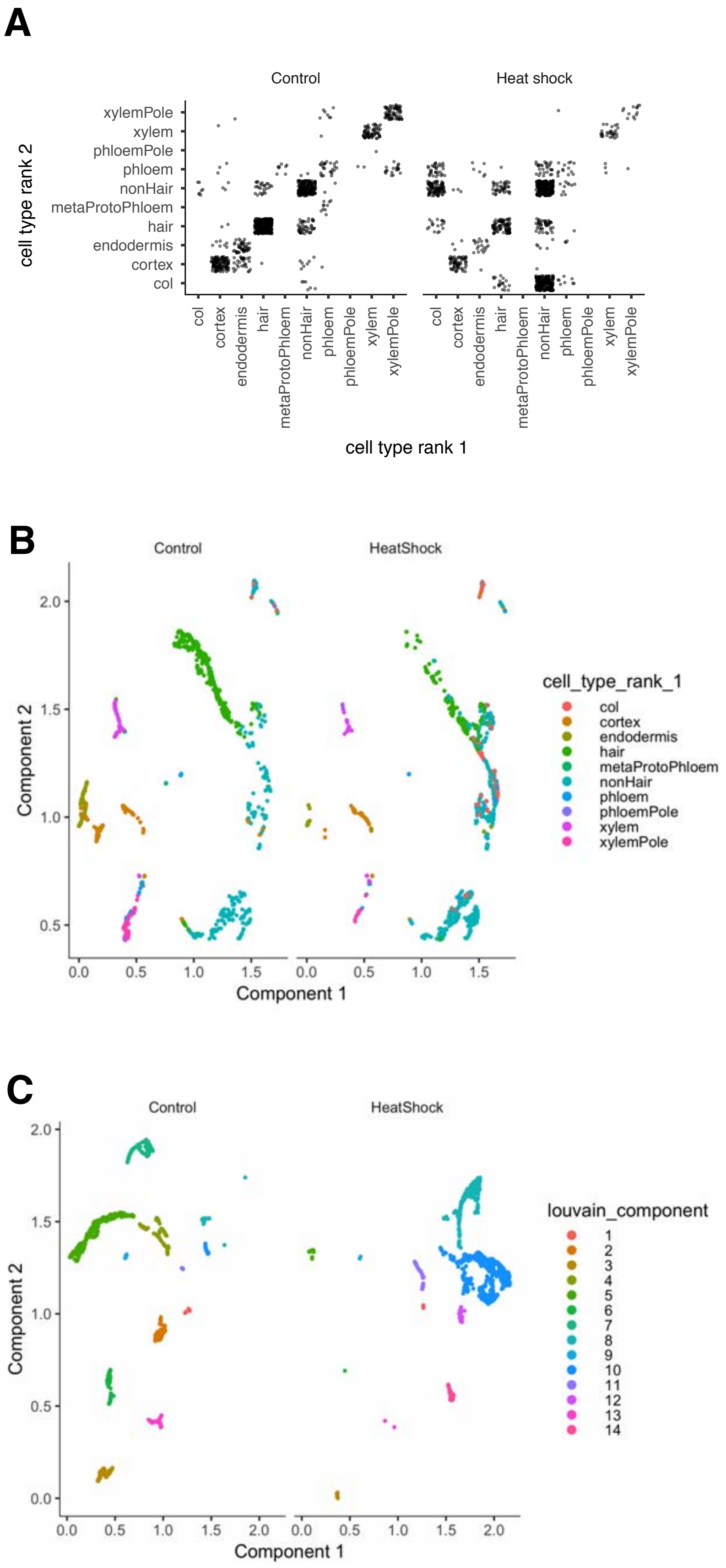


Supplemental Figure 11

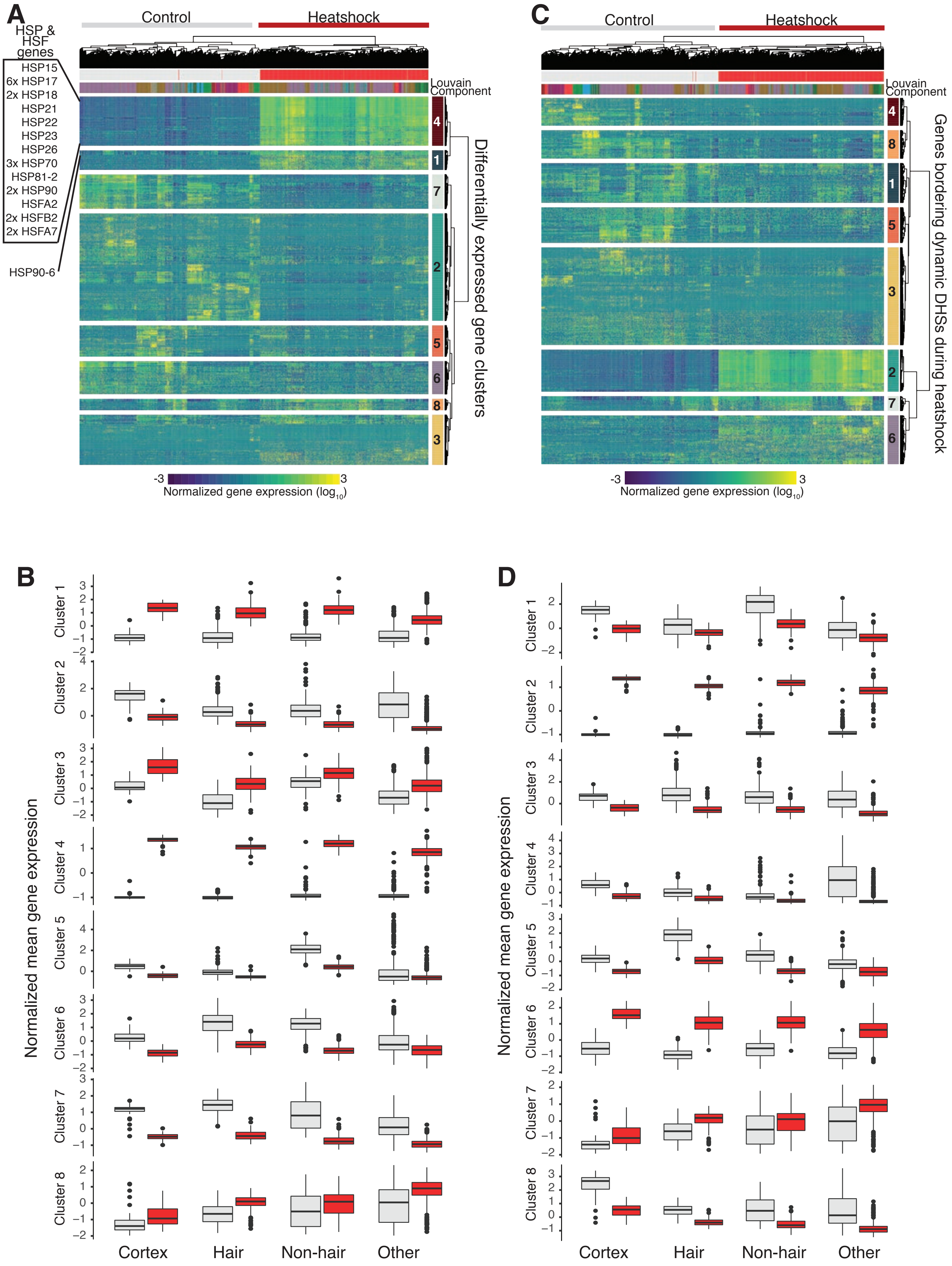

A
Control
Heatshock
HSP & HSF genes
HSP15
6x HSP17
2x HSP18
HSP21
HSP22
HSP23
HSP26
3x HSP70
HSP81-2
2x HSP90
HSFA2
2x HSFB2
2x HSFA7
HSP90-6
Louvain Component
Differentially expressed gene clusters
-3
3
Normalized gene expression (log10)
C
Control
Heatshock
Louvain Component
Genes bordering dynamic DHSs during heatshock
Normalized gene expression (log10)
B
Normalized mean gene expression
Cluster 1
Cluster 2
Cluster 3
Cluster 4
Cluster 5
Cluster 6
Cluster 7
Cluster 8
Cortex
Hair
Non-hair
Other
D
Normalized mean gene expression
Cortex
Hair
Non-hair
Other

**Supplemental Table 1.** Bulk RNA-seq comparisons to single cell RNA-seq

| Bulk GFP Line | Single Cell Group | Pearson Correlation | Spearman's Rank Correlation |
|---|---|---|---|
| **WOL** | WOL expressing cells | 0.45 | 0.66 |
| **WOL** | Stele | 0.51 | 0.68 |
| **WER** | WER expressing cells | 0.59 | 0.75 |
| **WER** | Non-hair | 0.54 | 0.72 |
| **SCR** | SCR expressing cells | 0.53 | 0.72 |
| **SCR** | Endodermis | 0.50 | 0.71 |
| **S4** | Stele | 0.37 | 0.57 |
| **S32** | Stele | 0.51 | 0.69 |
| **S18** | Stele | 0.66 | 0.80 |
| **Pet111** | Non-hair | 0.52 | 0.71 |
| **S17** | Stele | 0.66 | 0.80 |
| **GL2** | GL2 expressing cells | 0.58 | 0.74 |
| **GL2** | Non-hair | 0.62 | 0.78 |
| **COR** | Cortex | 0.74 | 0.86 |
| **E30** | Endodermis | 0.26 | 0.48 |
| **COBL9** | COBL9 expressing cells | 0.75 | 0.87 |
| **COBL9** | Hair | 0.74 | 0.87 |
| **APL** | Stele | 0.68 | 0.82 |
| **CO2** | Cortex | 0.3 | 0.55 |
| **Maturation** | Whole root | 0.49 | 0.70 |
| **Elongation** | Whole root | 0.7 | 0.83 |
| **Meristimatic** | Whole root | 0.21 | 0.42 |
| **Whole root** | Whole root | 0.52 | 0.71 |

**Supplemental Table 2.** Number of cells in the control vs. heat shock analysis

| Annotation / Louvain Component | Treatment ID | Number of Cells |
|---|---|---|
| Columella | Control | 7 |
| Columella | Heat Shock | 153 |
| Cortex | Control | 149 |
| Cortex | Heat Shock | 71 |
| Endodermis | Control | 77 |
| Endodermis | Heat Shock | 26 |
| Hair | Control | 371 |
| Hair | Heat Shock | 151 |
| MetaProtoPhloem | Control | 8 |
| MetaProtoPhloem | Heat Shock | 0 |
| NonHair | Control | 256 |
| NonHair | Heat Shock | 524 |
| Phloem | Control | 46 |
| Phloem | Heat Shock | 37 |
| PhloemPole | Control | 1 |
| PhloemPole | Heat Shock | 0 |
| Xylem | Control | 70 |
| Xylem | Heat Shock | 33 |
| XylemPole | Control | 91 |
| XylemPole | Heat Shock | 14 |
| Louvain Component 1 | Control | 24 |
| Louvain Component 1 | Heat Shock | 5 |
| Louvain Component 2 | Control | 400 |
| Louvain Component 2 | Heat Shock | 415 |
| Louvain Component 3 | Control | 160 |
| Louvain Component 3 | Heat Shock | 283 |
| Louvain Component 4 | Control | 97 |
| Louvain Component 4 | Heat Shock | 18 |
| Louvain Component 5 | Control | 120 |
| Louvain Component 5 | Heat Shock | 30 |
| Louvain Component 6 | Control | 77 |
| Louvain Component 6 | Heat Shock | 26 |
| Louvain Component 7 | Control | 60 |
| Louvain Component 7 | Heat Shock | 2 |
| Louvain Component 8 | Control | 41 |
| Louvain Component 8 | Heat Shock | 62 |
| Louvain Component 9 | Control | 16 |
| Louvain Component 9 | Heat Shock | 20 |
| Louvain Component 10 | Control | 43 |
| Louvain Component 10 | Heat Shock | 132 |
| Louvain Component 11 | Control | 38 |
| Louvain Component 11 | Heat Shock | 16 |
